# Gait-level analysis of mouse open field behavior using deep learning-based pose estimation

**DOI:** 10.1101/2020.12.29.424780

**Authors:** Keith Sheppard, Justin Gardin, Gautam S Sabnis, Asaf Peer, Megan Darrell, Sean Deats, Brian Geuther, Cathleen M. Lutz, Vivek Kumar

## Abstract

Gait and whole body posture are sensitive measures of the proper functioning of numerous neural circuits, and are often perturbed in many neurological, neuromuscular, and neuropsychiatric illnesses. Rodents provide a tractable model for elucidating disease mechanisms and interventions, however, studying gait and whole body posture in rodent models requires specialized methods and remains challenging. Here, we develop a simple assay that allows adoption of the commonly used open field apparatus for gait and whole body posture analysis. We leverage modern neural networks to abstract a mouse into keypoints and extract gait and whole body coordination metrics of the animal. Gait-level analysis allows us to detect every step of the animal’s movement and provides high resolution information about the animal’s behavior. We quantitate gait and whole body posture with high precision and accuracy across 62 highly visually diverse strains of mice. We apply our approach to characterize four genetic mutants with known gait deficits. In extended analysis, we demonstrate that multiple autism spectrum disorder (ASD) models show gait and posture deficits, implying this is a general feature of ASD. We conduct a large strain survey of 1898 mice, and find that gait and whole body posture measures are highly heritable in the laboratory mouse, and fall into three classes. Furthermore, the reference mouse strain, C57BL/6J, has a distinctly different gait and posture compared to other standard laboratory and wild-derived strains. We conduct a genome wide association study (GWAS) to define the genetic architecture of mouse movement in the open field. In sum, we describe a simple, sensitive, accurate, scalable, and ethologically relevant method of mouse gait and whole body posture analysis for behavioral neurogenetics. These results provide one of the largest laboratory mouse gait-level data resources for the research community and show the utility of automated machine learning approaches for deriving biological insights.

## 2 Introduction

In humans, the ability to quantitate gait and posture at high precision and sensitivity allows determination of proper function of numerous neural and muscular systems [1, 2]. Many psychiatric, neurodegenerative, and neuromuscular illnesses are associated with alterations in gait and posture, including autism spectrum disorder, schizophrenia, bipolar disorder, and Alzheimer’s disease. [3–12]. This is because proper gait, balance, and posture are under the control of multiple nervous system processes [13, 14], which include critical sensory centers that process visual, vestibular, auditory, proprioceptive, and visceral inputs. Regions of the brain that directly control movement, such as the cerebellum, motor cortex, and brain stem, respond to cognitive and emotionality cues. Thus, gait and posture integrity reflects proper neural functioning of many neural systems in humans [13, 14]. In rodent models of human psychiatric conditions, to date there has not been conclusive demonstrated utility of gait and posture metrics as in humans. This is not necessarily because the models fail to faithfully recapitulate human phenotypes, but rather that we have lacked readily-implementable technology with sufficient accuracy and scale to detect gait and posture differences between different mouse strains. The end result of this technological limitation is two-fold: rodent gait analysis remains tedious and often carried out only in expert labs with highly specialized equipment; and the behavioral neurogenetics field broadly has not been able to fully leverage relevant mouse gait phenotypes to understand the human disease. More importantly, while gait and movement analysis in humans is already a sensitive measure of numerous psychiatric illnesses, gait analysis in mice is generally used to study strong overt phenotypes. Thus, the ability to measure gait and whole body posture in an accurate, sensitive, and scalable manner is expected to enhance the utility of existing models and also lead to the development of better models of psychiatric endophenotypes.

Analysis of human and animal movement, including gait, has a storied past [15]. Aristotle was the first to write a philosophical treatise on animal movement and gait using physical and metaphysical principles [16]. During the Renaissance, Borelli applied the laws of physics and biomechanics to muscles, tendons, and joints of the entire body to understand gait [17]. The first application of imaging technologies to the study of gait is credited to the work of Muybridge and Marey, who took sequential photographic images of humans and animals in motion in order to derive quantitative measurements of gait [18–20]. Modern animal gait analysis methods are credited to Hildebrandt, who in the 1970s classified gait based on quantified metrics [21]. He defined a gait cycle in terms of contact of limb to the ground (stance and swing phases). Fundamentally, this concept has not changed over the past 40 years: while current methods of mouse gait analysis have increased efficiency of the imaging approaches of Muybridge and Marey, they are fundamentally still based on the timing of limbs contacting the ground. This is in contrast to human gait and posture analysis, which, since the time of Borelli, has focused on body posture, and is akin to the quantitation of whole body movement rather than just contact with the ground [22]. This difference between mouse and human is probably due in part to the difficulty in automatically estimating the posture of rodents, which appear as deformable objects due to their fur which obscures joint positions. In addition, unlike humans, parts of mice cannot be easily marked with wearables for localization. In rodents, recent methods have made progress by incorporation of speed in gait analysis [23, 24] and determination of whole body coordination [25, 26]. However, current common still require specialized equipment and force the animal to walk in a fixed direction in a treadmill or a narrow corridor for proper imaging and accurate determination of limb position [27]. This is highly unnatural, and animals often require training to perform this behavior properly [28–31], limiting the use of this type of assay in correlating to human gait. Imaging from the side leads to perspective hurdles, which are overcome by limiting the movement of the animal to one depth field. Furthermore, as the animal defecates and urinates, or when bedding is present, the resulting occlusion makes long term monitoring from this perspective impractical. Indeed, ethologically relevant gait data in which animals can move freely often produce results that differ from treadmill-based assays [32]. Furthermore, commercial treadmill-or corridor-based systems for gait analysis often produce a plethora of measures that show differing results with same animal models [27, 33]. The exact causes of these disparities are challenging to determine with closed, proprietary systems. Thus, we currently lack a relatively easily and broadly implementable tool to measure gait in free-moving animals at scale.

The open field assay is one of the oldest and most commonly used assays in behavioral neurogenetics [34, 35]. In rodents, it has classically been used to measure endophenotypes associated with emotionality, such as hyperactivity, anxiety, exploration, and habituation in rodents [36]. For video based open field assays, rich and complex behaviors of animal movement are often abstracted to a simple point in order to extract behavioral measures [37]. This oversimplified abstraction is necessary mainly due to technological limitations that have prohibited accurate extraction of complex poses from video data [38]. New technology has started to overcome this limitation [39–41] and has enabled a new era of animal behavior analysis. Gait, an important indicator of neural function, is not typically analyzed in the open field mainly due to the technical difficulty of determining limb position when animals are moving freely [27]. The ability to combine open field measures with gait and posture analysis would offer key insights into neural and genetic regulation of animal behavior in an ethologically relevant manner. Here, we leverage modern neural network methods to carry out mouse gait and posture analysis in the open field. We develop and apply a system to measure gait and whole body posture parameters from a top-down perspective that is invariant to the high level of visual diversity seen in the mouse, including coat color, fur differences, and size differences [42]. Altogether, we provide a methodology that is simple, sensitive, accurate, and scalable and can detect previously undescribed differences in gait and posture in mouse models of psychiatric illnesses. This method is a community resource for mouse movement in the open field that should commoditize gait analysis for behavioral neurogenetics.

## 3 Results

Our approach to gait and posture analysis is composed of several modular components. At the base of our toolkit is a deep convolutional neural network that has been trained to perform pose estimation on top-down video of an open field. This network provides twelve two-dimensional markers of mouse anatomical location, or “keypoints”, for each frame of video describing the pose of the mouse at each time point. We have also developed downstream components that are capable of processing the time series of poses and identifying intervals that represent individual strides. These strides form the basis of almost all of the phenotypic and statistical analyses that follow. We can extract several important gait metrics on a per-stride basis because we have pose information for each stride interval (see Table 1 for a list of metrics). This gives us significant power to perform statistical analysis on stride metrics as well as allowing us to aggregate large amounts of data in order to provide consensus views of the structure of mouse gait.

**Table 1:** Gait metrics definitions.

| Measure | Definition of Measure | Units |
| --- | --- | --- |
| Angular Velocity | The current angle of a mouse is determined by the vector connecting the mouse's base of tail to its base of neck. The first derivative of this value gives us angular velocity. For strides, angular velocity is averaged over the duration of the stride. | degrees/sec |
| Stride Speed | The speed of a mouse is determined by tracking the movement speed of the base of tail keypoint. Stride speed is the average speed for all frames over the duration of a stride. We shorten "Stride Speed" to "Speed" in some figure labels for compactness. | cm/sec |
| Limb Duty Factor | The stance time of a paw (the amount of time that the paw is in contact with the ground) divided by the full stride time. Duty factor is calculated for each of the hind paws and averaged. | None |
| Temporal Symmetry | Where $l$ is the left hind paw's duty factor and $r$ is the right hind paw's duty factor temporal symmetry is calculated as $(l - r)/(l + r)$ | None |
| Step Length | The distance that the right hind paw travels past the previous left hind paw strike | cm |
| Step Width | The averaged lateral distance separating hind paws. This is calculated as length of the shortest line segment that connects the right hind paw strike to the line that connects the left hind paw's toe-off location to its subsequent foot strike position. | cm |
| Stride Length | The full distance that the left hind paw travels for a stride, from toe-off to foot-strike. | cm |
| Lateral Displacement of Nose | In order to calculate lateral displacement, we first calculate the mouse's displacement vector for a stride. We then measure the nose's perpendicular distance from this vector for each frame of a stride. We now subtract the minimum distance from the maximum and divide by the mouse's body length so that the displacement measured in larger mice will be comparable to the distance measured in smaller mice. | None |
| Lateral Displacement of Base Of Tail | Calculated using the same approach which is applied to the Nose Lateral Displacement except that we are using the Base of Tail keypoint. | None |
| Lateral Displacement of Tip of Tail | Calculated using the same approach which is applied to the Nose Lateral Displacement except that we are using the Tip of Tail keypoint. | None |
| Nose Lateral Displacement Phase Offset | First, the lateral displacement is calculated for each frame of a stride as described for Nose Lateral Displacement above. We then perform a cubic spline interpolation in order to generate a smooth curve for displacement. We then determine the point in time where maximum displacement occurs. Note that because of cubic interpolation this can occur at time points between frames. | Percent Stride Cycle |
| Base of Tail Displacement Phase Offset | Calculated using the same approach which is applied to the Nose Lateral Displacement Phase Offset except that we are using the Base of Tail keypoint. | Percent Stride Cycle |
| Tip of Tail Displacement Phase Offset | Calculated using the same approach which is applied to the Nose Lateral Displacement Phase Offset except that we are using the Tip of Tail keypoint. | Percent Stride Cycle |

### 3.1 Pose Estimation

Pose estimation locates the 2D coordinates of a pre-defined set of keypoints in an image or video, and is the foundation of our method for quantifying and analyzing gait. The selected pose keypoints are either visually salient, such as ears or nose, or capture important information for understanding pose, such as limb joints or paws. We thus selected twelve keypoints to capture mouse pose: nose, left ear, right ear, base of neck, left forepaw, right forepaw, mid spine, left hind paw, right hind paw, base of tail, mid tail and tip of tail (Figure 1B).

**Figure 1:**
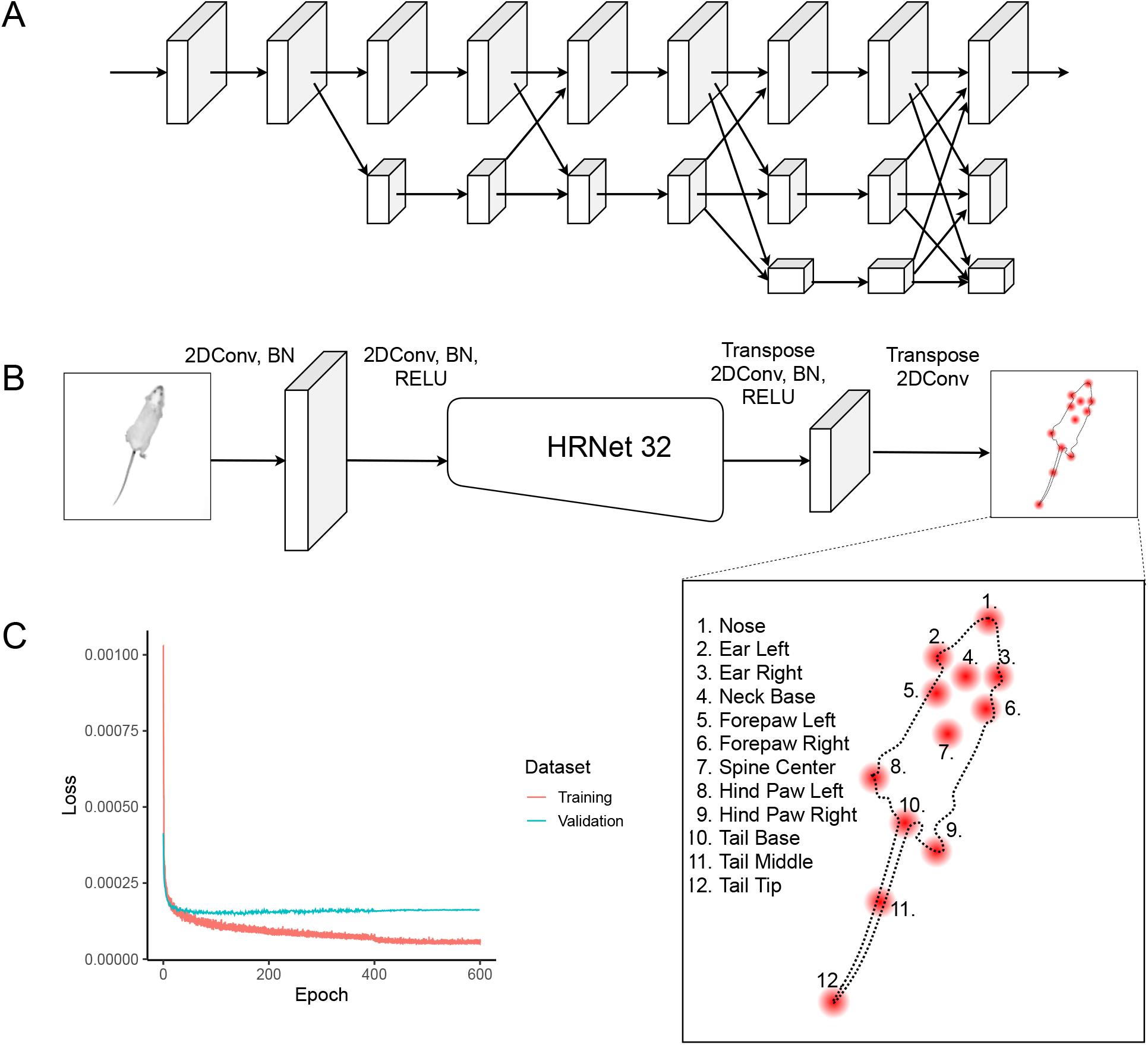
Deep convolutional neural network for pose estimation. (A) the HRNet-W32 neural network architecture for performing pose estimation. (B) The inference pipeline which sends video frames into the HRNet and generates twelve keypoint heatmaps as output. (C) Training loss curves show network convergence without overfitting.

Much effort has been spent developing and refining pose estimation techniques for human pose [43, 44]. Traditional approaches to pose estimation relied on techniques such as the use of local body part detectors and modeling of skeletal articulation. These approaches were limited in their ability to overcome complicating factors such as complex configurations and body part occlusion. The first major paper to address these shortcomings by developing a deep neural network for pose estimation was the DeepPose [45]. DeepPose was able to demonstrate improvements on the state-of-the-art performance for pose estimation using several benchmarks. After the publication of DeepPose, the majority of successful work on pose estimation leveraged deep convolutional neural network architectures. Some prominent examples include: DeeperCut [46], Stacked Hourglass Networks [47] and Deep High-Resolution architecture (HRNet) [48]. This is a rapidly evolving field and architectural improvements are frequently released and tracked by several leaderboards [49]. Given this choice of high performance pose estimation architectures developed for human pose estimation, it made sense to leverage this prior work for our rodent pose estimation problem rather than attempt to develop our own architecture. There were several important considerations we based our pose estimation architecture selection upon.

- High accuracy and precision for pose inference: our gait inference method is sensitive to errors in pose estimation so we want to reduce those errors as much as possible
- Speed of inference: should be able to infer at or near real time speeds (30 fps) on a modern high end GPU
- Fixed scale inference: because all of our images are at fixed scale, approaches that are designed to work at multiple scales waste network capacity and inference time.
- Available open source implementation
- Modularity of architecture in order to facilitate potential future upgrades.

Based on these criteria we selected the HRNet architecture [48] for our network and modified it for our experimental setup. The main differentiator of this architecture is that it maintains high resolution features throughout the network stack, thereby preserving spatial precision (Figure 1A). HRNet shows highly competitive performance in terms of both GPU efficiency and pose accuracy. The interface is also highly modular and is expected to allow for relatively simple network upgrades if needed. HRNet has been compared for performance against other state of art networks including DeeperCut, on which DeepLabCut is based [39]. HRNet outperforms these networks for accuracy and compute. For instance, HRNet-32 achieves a Percentage of Correct Key-points (PCK .5) value of 92.3 compared to 88.5 for DeepLab. In addition HRNet has far fewer parameters (28.5M vs. 42.6M) and uses less compute (9.5 vs 41.2 GFLOPS) (Table 3 and 4 of Sun et. al. [48]). We used the smaller HRNet-W32 architecture rather than HRNet-W48 because it was shown to provide significant speed and memory improvements for only a small reduction in accuracy. We added two 5×5 transpose convolutions to the head of the network to match the heatmap output resolution with the resolution of the video input (Figure 1B). Because all of our experiments have a single mouse in an open field, we do not need to rely on object detection for instancing. We thus eliminated this step from our inference algorithm, which also leads to clear runtime performance benefits. Instead of performing pose estimation after object detection, we use the full resolution pose keypoint heatmaps to infer the posture of a single mouse at every frame. This means that for each 480×480 frame of video we generate 12 480×480 heatmaps (one heatmap per keypoint). The maximum value in each heatmap represents the highest confidence location for each respective point. Thus, after taking the argmax of each of the 12 heatmaps we have 12 (x, y) coordinates.

In order to train our network, we need to select a loss function and an optimization algorithm. For loss, we borrow the approach used in the original HRNet description [48]. For each keypoint label, we generate a 2D gaussian distribution centered on the respective keypoint. We then compare the output of the network with our keypoint-centered Gaussian and calculate loss as the mean squared difference between our labeled keypoint Gaussian and the heatmap generated by our network. We train our network using the ADAM optimization algorithm which is a variant of stochastic gradient descent [50]. Figure 1C shows that the validation loss converges rapidly. We intentionally generated labels that represent a wide diversity of mouse appearances, including variation in coat color, body length and obesity to ensure that the resulting network operates robustly across these differences. We manually labeled 8,910 frames across these diverse strains for training (see Methods). The resulting network is able to track dozens of mouse strains with varying body size, shape and coat color (Video S1) [42]. We calculate the accuracy of our neural network and experimental configuration over two datasets: a set of 1000 images with results for 200 white mice and 200 dark mice broken out (Table S1) as well as a set of 120 images containing twenty images each from a set of six visually diverse mouse strains (Figure S2). We provide these metrics in pixel and centimeter distance units.

### 3.2 Stride Inference

Our approach to detecting stride intervals is based on the cyclic structure of gait as described by Hildebrand (Figure 2A, B) [21, 51]. During a stride cycle, each of the paws has a stance phase and a swing phase [27]. During the stance phase, the mouse’s paw is supporting the weight of the mouse and is in static contact with the ground. During the swing phase, the paw is moving forward and is not supporting the mouse’s weight. Following Hildebrand, we refer to the transition from stance phase to swing phase as the toe-off event and the transition from swing phase to stance phase as the foot-strike event.

**Figure 2:**
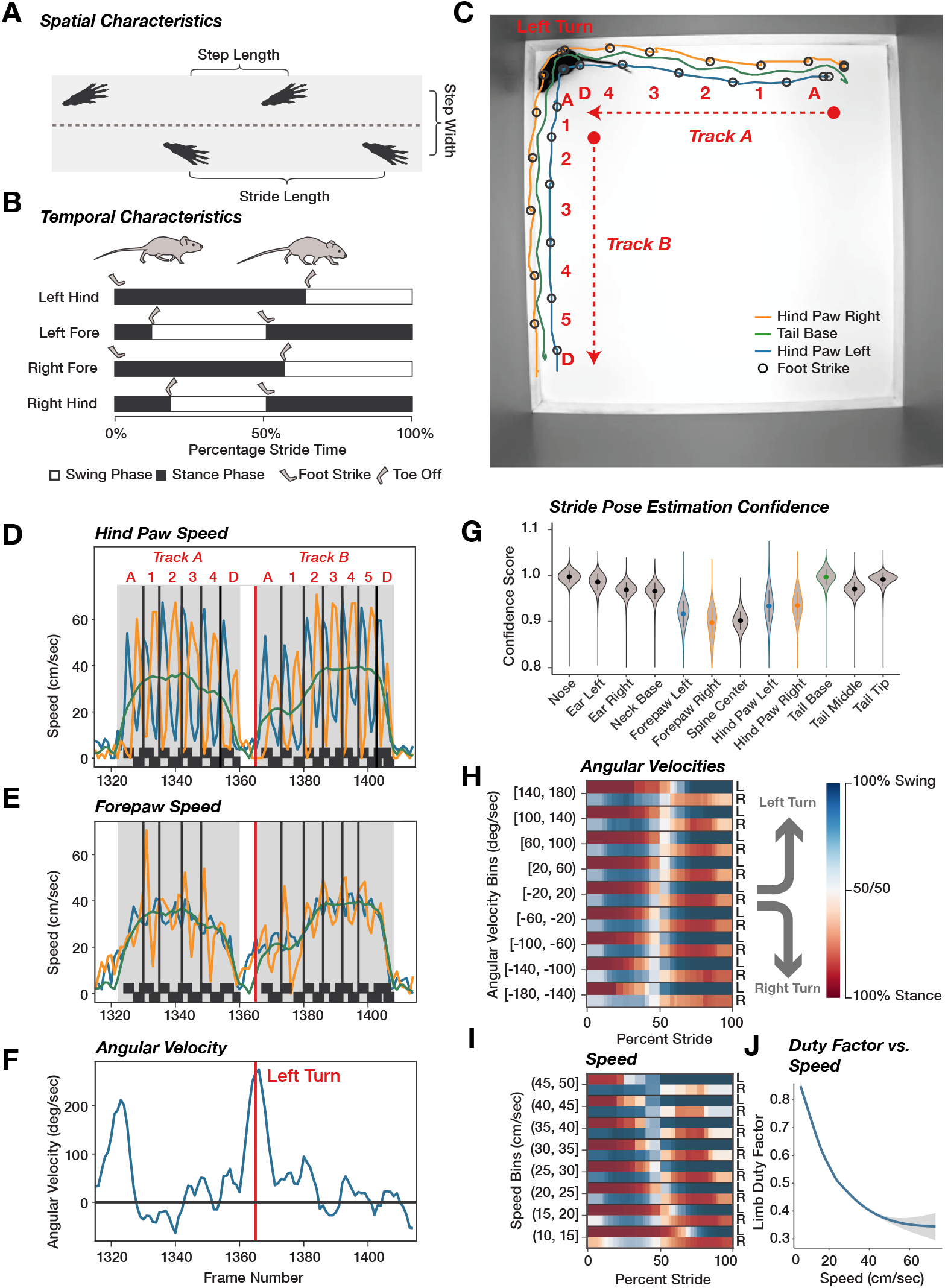
Extraction of gait metrics from video pose estimation. Spatial and temporal characteristics of gait (A, B, adapted from [27]). (A) An illustration showing how we derive three spatial stride metrics from hind paw foot strike positions: step length, step width and stride length. (B) All metrics shown in this Hildebrand plot have percent stride time for units. We see here the relationship between foot strike and toe off events with the stance and swing phases of stride. (C) A single frame of input video with hind paw tracks plotted fifty frames in the past and fifty frames in the future. The location of hind foot strike events is indicated with black circles and paths are shown (left hind paw (blue), right hind paw (orange), and base of tail (green)) for two sequences (track A and B). (D-F) Three plots showing different aspects of the mouse’s movement over the same one hundred frame interval (Video S2). The centered red vertical line indicates the current frame (displayed in panel C). The top plot shows three lines indicating speed of the left hind paw (blue), the right hind paw (orange) and the base of tail (green). The vertical black lines in the plot indicate the inferred start frame of each stride. (G) The distribution of confidence values for each of the 12 keypoints we estimate. (H) Aggregate view of Hildebrand plot for hind paws binned according to angular velocity (left (L) and right (R)) shows changes in strike duration based on direction of turning. (I) Similar to panel (H) except binned by increasing stride speed and a fixed angular velocity (−20 to 20 deg/sec). (J) Limb duty factor changes as a function of stride speed. Data for panels (H), (I) and (J) are derived from 15,667 strides from 31 C57BL/6NJ animals.

In order to calculate stride intervals, we determine stance and swing phases for the hind paws. We calculate paw speed and infer that a paw is in stance phase when the speed falls below a threshold and that it is in swing phase when it exceeds that threshold (Figure 2C, D, E, F, Video S2). We can now determine that foot strike events occur at the transition frame from swing phase to stance phase (Figure 2C). We define the left hind foot strike as the event that separates stride cycles. An example of the relationship between paw speed and foot strike events is shown in Figure 2D for hind paws. We find clean, high-amplitude oscillations of the hind paws, but not forepaws, as shown in Figure 2E. This difference in inference quality between the forepaws and hind paws is likely due to the fact that forepaws are occluded more often than hind paws from the top-down view and are therefore more difficult to accurately locate. We observe a corresponding decrease in confidence of forepaw inferences as show in Figure 2G. For this reason, we exclude forepaws from consideration when deriving stride intervals and focus instead on hind paws. We also perform a significant amount of filtering on strides to remove spurious or low quality stride cycles from our dataset (Figure 2G). Criteria for removing strides include: low confidence or physiologically unrealistic pose estimates, missing right hind paw strike event, and insufficient overall body speed of mouse which is any speed under 10 cm/sec. Panel G of Figure 2 shows the distribution of confidences for each keypoint. Our filtering method uses 0.3 as a confidence threshold. Very high confidence keypoints are close to 1.0. We always remove the first and last strides in a continuous sequence of strides to avoid starting and stopping behaviors from adding noise to our stride data (Figure 2C, D, labeled A and D, in Track A and B). This means that a sequence of seven strides will result in at most five strides being used for analysis. The distribution of keypoint confidence varies by keypoint type (Figure 2G). Keypoints which tend to be occluded in a top-down view such as fore paws have confidence distributions shifted down compared to other keypoints. We also see that keypoints that are not visually salient, such as the spine center, will have lower confidence since they are more difficult to locate precisely. Finally, we also calculate an instantaneous angular velocity which allows us to determine the turning direction of each stride (Figure 2F). The angular velocity is calculated by taking the first derivative of the angle formed by the line that connects the base of the mouse’s tail to the base of its neck. In sum, this approach allows us to identify individual high quality strides of a mouse in the open field.

In order to validate that our gait quantitation is functioning properly, we analyzed data from a commonly used inbred strain, C57BL/6NJ. We calculated percent of stance and swing from 15,667 strides from 31 animals using approximately 1-hour of open field video per mouse. We analyzed data from hind paws since these showed the highest amplitude oscillations during stance and swing (Figure 2D, E). We stratified the data into 9 angular velocity and 8 stride speed bins based on the tail base point (Figure 2H, I, respectively). As expected, we find increase in stance percent over a stride of the left hind paw when the animal is turning left. Reciprocally, when the animal is turning right, the stance percent of the right hind paw is increased (Figure 2H). We then analyzed strides in the central angular velocity bin (−20 to 20 deg/sec) to determine if stance percent during a stride cycle decreases as the speed of the stride increases. We find the stance time decreases as the stride speed increases (Figure 2I). We generated the same plots for five other mouse strains and see similar results for all five (Figure S1). We calculated a duty factor for the hind paws to quantitate this relationship with stride speed (Figure 2J). In sum, we conclude that our methods are able to quantitatively and accurately extract strides from these open field videos from a top-down perspective.

After the stride intervals have been determined, we use frame poses in conjunction with stance and swing phase intervals to derive several stride metrics as defined in Table 1. We are able to extract most relevant spatiotemporal metrics from the hind paws, which serve as the primary data source for our statistical analyses [27].

### 3.3 Whole body posture estimation during gait cycle

Our top-down videos allow us to determine the relative position of the spine with 6 keypoints (nose, neck base, spine center, tail base, tail middle, and tail tip). With these, we extracted the whole body pose during a stride cycle, similar to previous work which carried this out with nose and tail pose only [25]. We used three points (nose, base of tail, and tip of tail) to capture the lateral movement during a stride cycle (Figure 3A, B, C, Video S3). These measures are circular, with opposite phases of the nose and the tip of tail. For display, we use C57BL/6J (Video S5, S6) and NOR/LtJ (Video S4, S6) which have different tip of tail phases during a stride cycle. We are able to extract these phase plots for each stride (Figure 3D, E, Video S3, S4, S5, S6). Since we have several hours of video across each strain, we are able to extract thousands of strides enabling high level of sensitivity. We can combine these at one stride speed and angular velocity bin where we constrain the speed range from 20 to 25 cm/sec and angular velocity from -20 to 20 deg/sec to determine a consensus stride phase plot for each animal and strain (Figure 3F, G). Finally, we compared these phase plots between several strains and find striking diversity among whole body posture during the gait cycle (Figure 3H,I). The diversity in whole body coordination across mouse strains is evident and implies high heritability of this phenotype.

**Figure 3:**
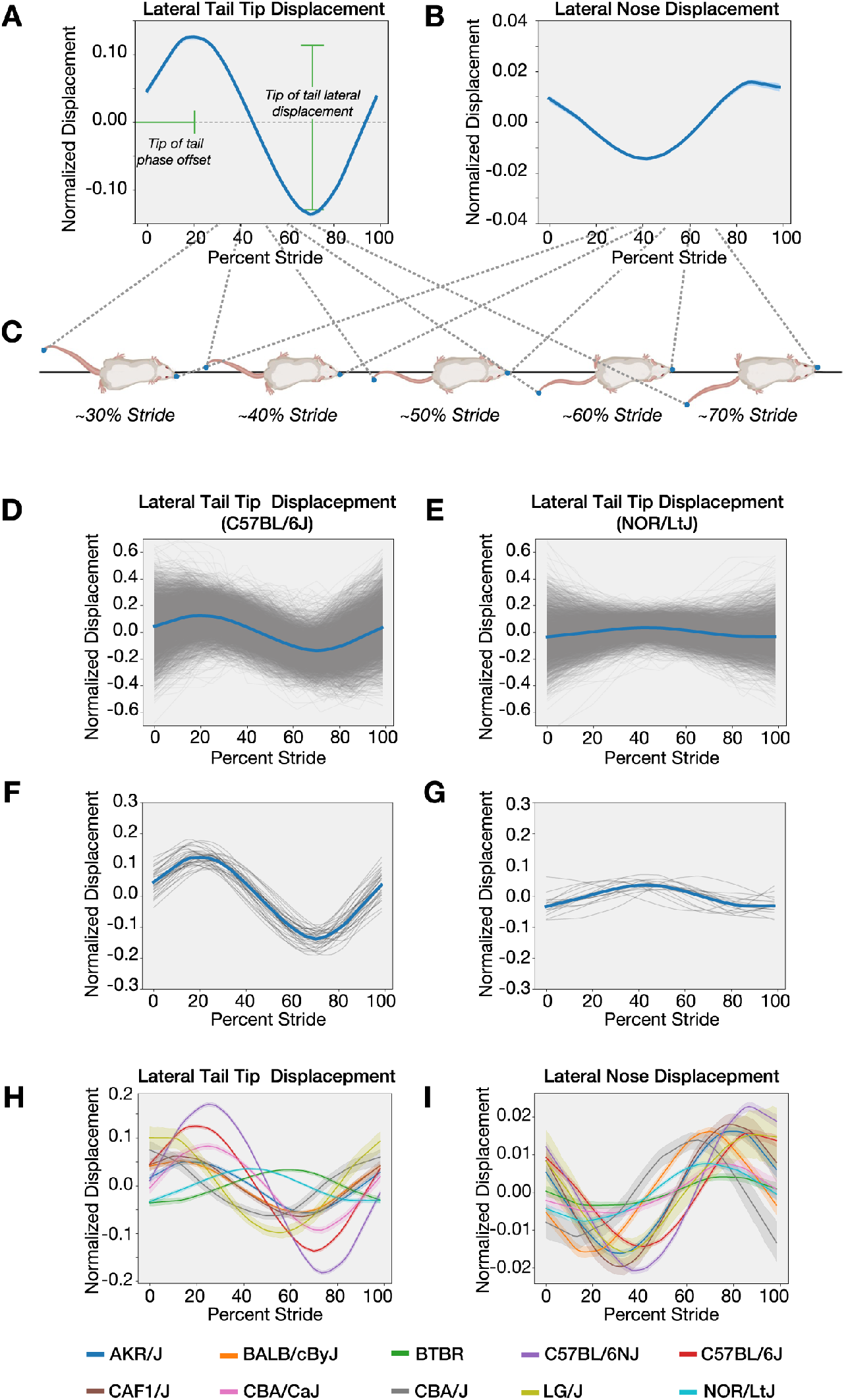
Extraction of cyclic whole body posture metrics during gait cycle. We measure lateral displacement of (A) the tail tip and (B) the nose. Positive values are to the animal’s left and negative values are to its right. We label this “Normalized Displacement” because displacement values are divided by the respective animal’s body length. We do this so that differences in amplitude cannot simply be attributed to animal size. We can also average displacement across many strides within a cohort to form a consensus view such as (D) C57BL/6J vs. (E) NOR/LtJ or we can average many strides within individuals: (F) C57BL/6J vs. (G) NOR/LtJ. For tail (H) and nose (I) we see the diversity of lateral displacement between a set of strains selected from our strain survey. The translucent bands for these two plots represent the 95% confidence interval of the mean for each respective strain.

Several of our metrics relate to the cyclic lateral displacement we observe in pose keypoints (Figure 3). Our measures of lateral displacement are defined as an orthogonal offset from the relevant stride displacement vector. We define the displacement vector as the line connecting the mouse’s center of spine on the first frame of a stride to the mouse’s center of spine on the last frame of stride. We calculate this offset at each frame of a stride and then perform a cubic interpolation in order to generate a smooth displacement curve. The phase offset of displacement is defined as the percent stride location where maximum displacement occurs on this smoothed curve. As an example, if we observe a value of 90 for phase offset it indicates that the peak lateral displacement occurs at the point where a stride cycle is 90% complete. The lateral displacement metric assigned to stride is the difference between maximum displacement value and minimum displacement value observed during a stride (Figure 3A). This analysis is sensitive and allows us to detect subtle, but highly significant differences in overall posture during a stride (Video S4, S5, S6). We used the previous classical spatiotemporal measures based on Hildebrand’s methods with the combined whole body posture metrics for our analysis. Because of the cyclic nature of phase offset metrics, care was taken to apply circular statistics to these metrics in our analysis. The other measures are analyzed using linear methods.

Next, we determined whether stride metrics changed depending on the location of the animal. For instance, animals displaying thigmotaxis are considered to be more anxious and in a lower state of arousal than those that are in the center [36]. In order to determine if these differing emotional states affect gait and whole body coordination metrics, we analyzed the strides based on the location at which they occur. We partitioned the each stride into center or periphery strides. To carry this out we trained a new neural network to detect corners of our open field. We defined the periphery as the outermost 10% of the matrix (Figure S7A blue vs. purple). We only analyzed strides in 20-25 cm/sec and 25-30 cm/sec speed bins with angular velocity in (−20,20) degrees/sec. Both groups contained an approximately equal number of strides for both strains (Figure S7B red vs. blue). Analysis of the gait and whole body coordination metrics showed no difference between the center and periphery (Figure S7C,D). Surprisingly, this analysis indicates that gait and whole body coordination measures do not change in response to location and by extension emotional state of the animal.

### 3.4 Statistical Analysis and genetic validation of gait measures

Following gait and posture extraction, we established a statistical framework for analysis of the data. In order to validate our methods, we phenotyped three mouse models that have previously been shown to have gait defects and are preclinical models of human diseases -Rett’s syndrome, Amyotrophic Lateral Sclerosis (ALS or Lou Gehrig’s Disease), and Down syndrome. The three models, *Mecp2* knockout, SOD1 G93A transgene, and *Ts65Dn Trisomic*, respectively, were tested with appropriate controls at two ages in a one hour open field assay (Table 2). Gait metrics are highly correlated with animal size and stride speed [23–26, 51] (Figure 2I, J). However in many cases changes in stride speed is a defining feature of gait change due to genetic or pharmacological perturbation. In addition, we have multiple repeated measurements that are collected for each subject (mouse) and each subject has a different number of strides giving rise to imbalanced data. Averaging over repeated strides, which yields one value per subject, can be misleading as it removes variation and introduces false confidence. At the same time, classical linear models do not discriminate between stable intra-subject variations and inter-subject fluctuations which severely bias the estimates. To address this, we used a linear mixed model (LMM) to dissociate within-subject variation from genotype-based variation between subjects [52, 53]. Specifically, in addition to the main effects such as animal size, genotype and age, a random effect that captures the intra-subject variation is included. Finally, we have multiple repeated measurements at two different ages giving rise to a nested hierarchical data structure. The models (M1, M2 M3) follow the standard LMM notation with (Genotype, BodyLength, Speed, TestAge) denoting the fixed effects and (MouseID/TestAge) (test age nested within the animal) denoting the random effect. In order to compare our results with previously published data that do not take animal size and sometimes stride speed into account, we statistically modeled our data with three models that only take age and body length (M1), age and stride speed (M2), age, stride speed, and body length (M3) as covariates (Figure 4 and S2).

**Table 2:** Control strains and official identifiers for gait mutants.

| Strain | JMCRS # | Genotype | Sex | Short Name | Control Strain |
| --- | --- | --- | --- | --- | --- |
| B6.129P2(C)-Mecp2<sup>tm1.1Bird</sup>/J | 003890 | HEMI | M | Mecp2 | Littermate WT |
| B6.129P2(C)-Mecp2<sup>tm1.1Bird</sup>/J | 003890 | HET | F | Mecp2 | Littermate WT |
| B6.Cg-Tg(SOD1*G93A)1Gur/J | 004435 | HEMI | M/F | Sod1 | Littermate WT |
| B6EiC3Sn.BLiA-Ts(17<sup>16</sup>)65Dn/DnJ | 005252 | Trisomic | M/F | Ts65Dn | B6EiC3Sn.BLiAF1/J |
| B6EiC3Sn.BLiAF1/J | 003647 | F1 | M/F |  | N/A |

**Table 3:**
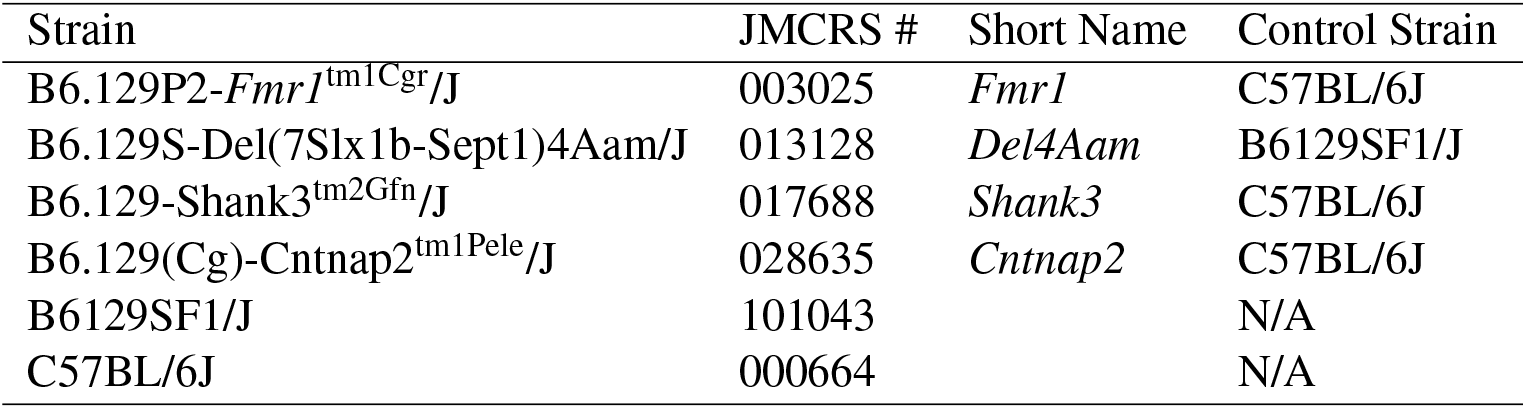
Control strains and official identifiers for autism mutants.

| Strain | JMCRS # | Short Name | Control Strain |
| --- | --- | --- | --- |
| B6.129P2-*Fmr1*<sup>tm1Cgr</sup>/J | 003025 | *Fmr1* | C57BL/6J |
| B6.129S-Del(7Slx1b-Sept1)4Aam/J | 013128 | *Del4Aam* | B6129SF1/J |
| B6.129-Shank3<sup>tm2Gfn</sup>/J | 017688 | *Shank3* | C57BL/6J |
| B6.129(Cg)-Cntnap2<sup>tm1Pele</sup>/J | 028635 | *Cntnap2* | C57BL/6J |
| B6129SF1/J | 101043 |  | N/A |
| C57BL/6J | 000664 |  | N/A |

**Table 4:**
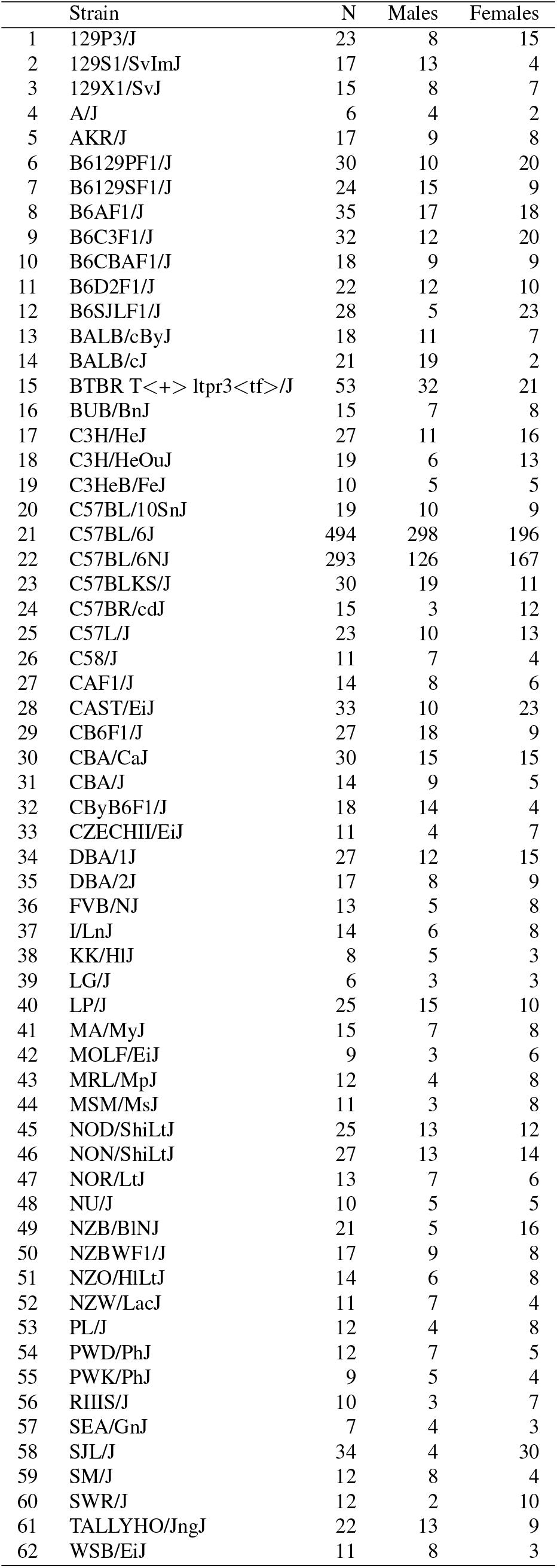
Summary data for strain counts in the strain survey

**Figure 4:**
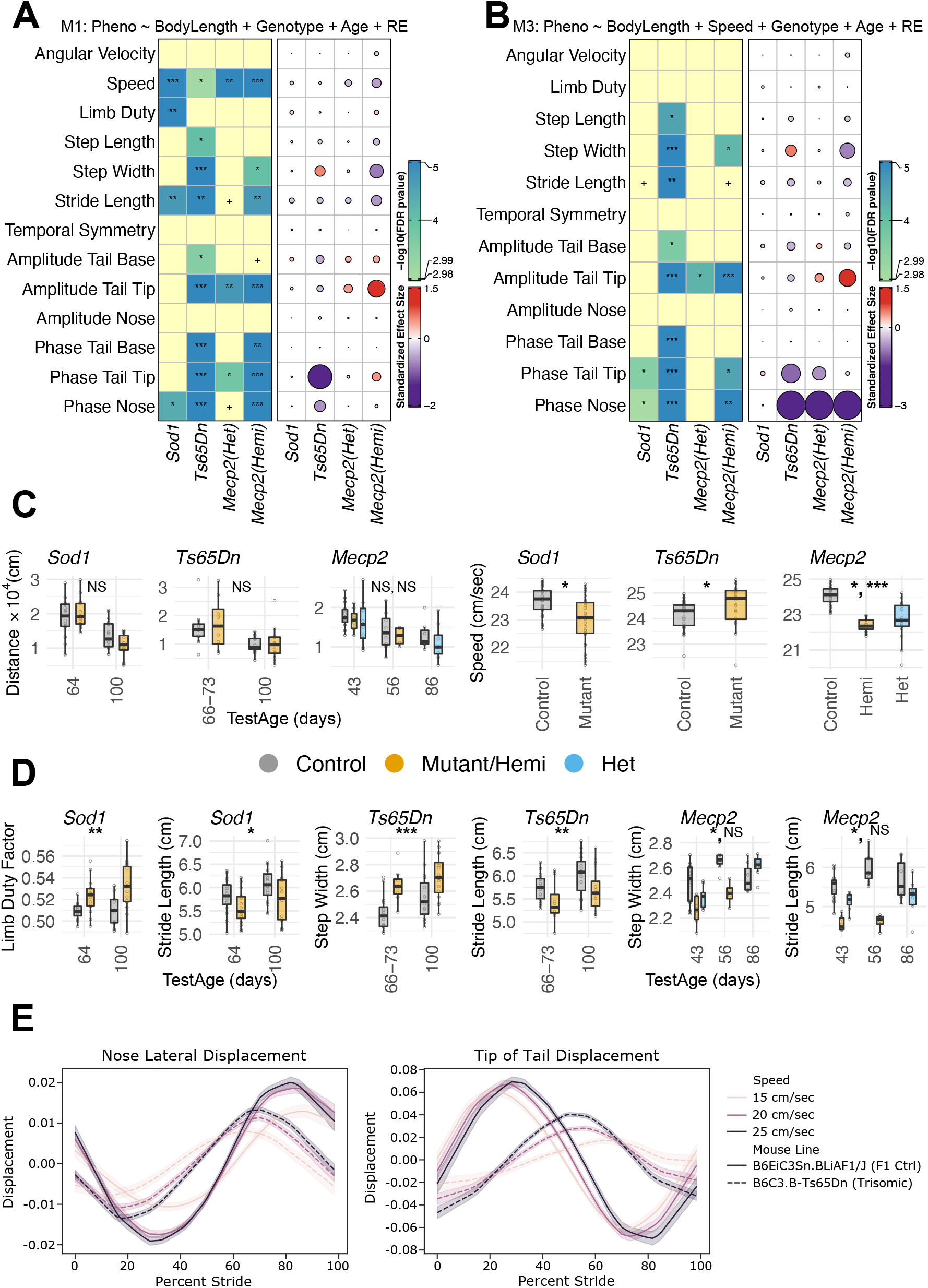
Genetic validation of the gait extraction methods. (A) We validate previous findings using M1, which adjusts only for body length, TestAge (Age), Genotype and random effects (RE). The LOD (*−*log_10_(q-value)) scores and effect sizes are shown in the left and right vertical blocks, respectively. In the left block, the number of ‘*★*’ and heat represent the strength of evidence against the null hypothesis of no genotype-based effect while + represents a suggestive effect. In the right block, the color (red for +ve and blue for -ve) and area of the circle (area *∝* size of the effect) represent the effect size’s direction and magnitude. (B) Same as (A), except that we validate previous findings using model M3, which adjusts for both body length, stride speed (speed), Genotype, TestAge (Age) and random effects (RE). (C) We plot Distance *×*10^4^ (cm), across test ages (x-axis), and (stride) Speed (cm/sec) across gait mutants (*Sod1, Ts65Dn, Mecp2*) for comparing mutants with controls. Each dot represents a tested animal. (D) We plot the most significant gait parameters from (A) for different gait mutants to compare mutants with controls across test ages (x-axis). (E) Lateral displacement of nose and tail tip for Ts65Dn strain. The solid lines represent the mean displacement of stride, while the translucent bands provide a 95% confidence interval for the mean.

M1 : Phenotype *∼* Genotype + TestAge + BodyLength + (1 | MouseID*/*TestAge)

M2 : Phenotype *∼* Genotype + TestAge + Speed + (1 | MouseID*/*TestAge)

M3 : Phenotype *∼* Genotype + TestAge + Speed + BodyLength + (1 | MouseID*/*TestAge)

In general, we use M1 to detect changes in stride speed and M3 for changes in gait parameters. We include data for M2 in supplement for comparison with previously published data. We do not include sex in our models as it is highly correlated with body length (measured using ANOVA and denoted by *η*, is strong for both SOD1 (*η* = 0.81) and *Ts65Dn* (*η* = 0.16 overall, *η* = 0.89 for controls, *η* = 0.61 for mutants). We analyze *Mecp2* males and females separately. We model the circular phase variables in Table 1 as a function of linear variables using a circular-linear regression model [54]. To adjust for linear variables such as body length and stride speed, we include them as covariates in the model (also see Methods). We report LOD scores, defined as -log(q-value) where q-value is the FDR adjusted p-value, and effect sizes in Figures 4, 5. For clarity, exact statistics are reported in detail in the supplementary tables S3, S4.

**Figure 5:**
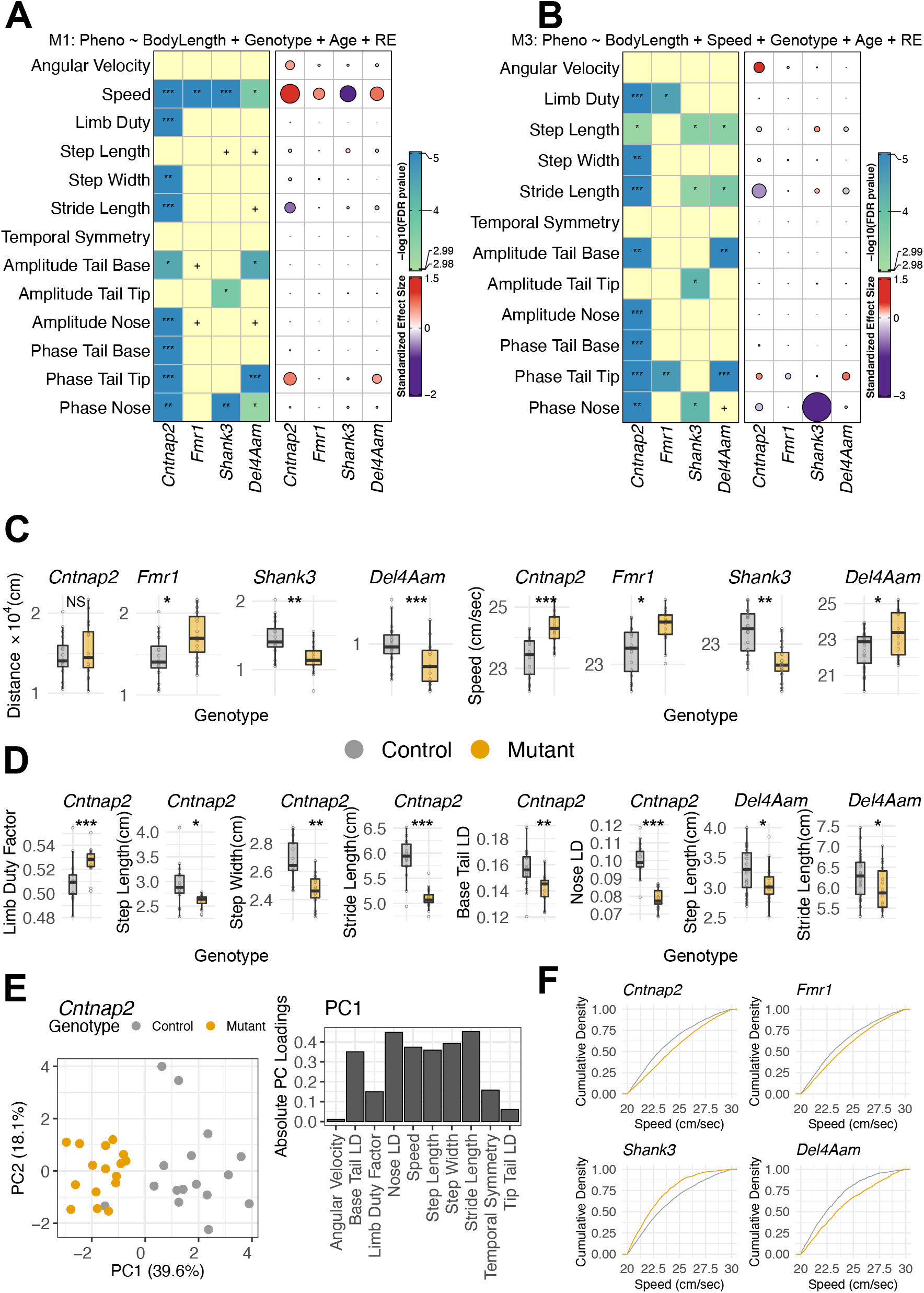
Characterization of gait and posture in mouse genetic models of autism. (A) We validate previous findings using model M1, which adjusts only for body length. The LOD scores and effect sizes are shown in the left and right vertical blocks, respectively. In the left block, the number of *\**s represents the strength of evidence against the null hypothesis of no genotype-based effect while + represents a suggestive effect. In the right block, the color (red for +ve and blue for -ve) and area of the circle (area *∝* size of the effect) represent the effect size’s direction and magnitude. (B) Same as (A), except that we validate previous findings using model M3, which adjusts for both body length and stride speed. (C) We plot Distance *×* 10^4^ (cm), across test ages (x-axis), and stride Speed (cm/sec) across autism mutants (*Cntnap2, Fmr1, Shank3, Del4Aam*) for comparing mutants with controls. Each dot represents a tested animal. (D) We plot the most significant gait parameters from (B) for different gait mutants to compare mutants with controls. (E) We perform PCA on z-score transformed gait data for *Cntnap2* mutants and their controls. We use the first two principal components to plot a 2D representation of the multidimensional gait space, which separates controls from the mutants. The dots represent individual animals. (F) We plot the stride speed cumulative distributions between mutants and controls across autism mutants.

#### 3.4.1 Validation using a Rett syndrome model

Rett syndrome, an inherited neurodevelopmental disorder, is caused by mutations in the X-linked *MECP2* gene [55]. We tested a commonly studied deletion of *Mecp2* that recapitulates many of the features of Rett syndrome, including reduced movement, abnormal gait, limb clasping, low birth weight, and lethality [56]. We tested hemizygous males (*n* = 8), heterozygous females (*n* = 8), and littermate controls (*n* = 8 of each sex) (Table 2). Null males are normal at birth and have an expected lifespan of about 50-60 days. They start to show age-dependent phenotypes by 3-8 weeks and lethality by 10 weeks. Heterozygous females have mild symptoms at a much older age [56]. We tested male mice twice at 43 and 56 days and females at 43 and 86 days.

Previous gait studies of this knockout did not take animal size, and in some cases, changes in stride speed into account. These studies have shown changes in stride length and stance width in an age-dependent manner in hemizygous males [57–59]. Recent analysis showed increased step width, reduced stride length, changes in stride time, step angle, and overlap distance [60]. *Mecp2* hemizygous males show 13% reduced body (Table 5, Figure S3C) [56] and progressive changes in movement speed that should be taken into account when modeling gait parameters. We limit our analysis to stride speeds between 20-30cm/s which allows us to reduce variation introduced by differences in speed and compare a model that includes body length instead of stride speed as a covariate (M1, Figure 4A) and one in which both body length and stride speed are included (M3, 4B). We placed the results from M2 into supplement for comparison with previously published data (Figure S3) and all statistics (M1, M2, M3) are reported in Table 5. Model M3 that includes both stride speed and body length showed a significant decrease in step width and suggestive difference in stride length, and robust differences in whole body coordination metrics (tail tip amplitude, phase of tail tip, and nose) (Figure 4B). We also note a decrease in total distance traveled in the open field, stride speed, stride length, and step width in the mutants after adjusting for body length (M1) (Figure 4C, D). Even though we limit the analysis to one angular and speed bin, we see differences in distribution of stride speed (Figure S2B). We observe very few significant differences in *Mecp2* heterozygous females that are consistent across all three models. All three models consistently find tail tip amplitude to be significantly higher suggesting more lateral movement in the females (Figure 4A,B and S2). In sum, these results demonstrate that we are able to accurately detect previously described differences in *Mecp2*. In addition, our whole body coordination metrics are able to detect differences that have not been previously described. All three models consistently find tail tip amplitude to be significantly higher suggesting more lateral movement in the females (Figure 4A,B and S2A).

**Table 5:** Summary data for body length and weight of animals in our experiments.

| Gait Mutants |  |  |  |  |  |  |
| --- | --- | --- | --- | --- | --- | --- |
| Strain | Sex | BodyLength (cm) |  | Correlation | BodyWeight (g) |  |
| | | Control | Mutant | $r$ | Control | Mutant |
| *Sod1* | M | 6.16 ± .15 | 6.23 ± .26 | 0.48 | 28.90 ± 2.69 | 26.41 ± 2.25 |
| | F | 5.53 ± .31 | 5.47 ± .32 | 0.76 | 21.74 ± 2.38 | 20.08 ± 1.84 |
| *Ts65Dn* | M | 5.90 ± .29 | 5.75 ± .37 | 0.73 | 30.46 ± 4.16 | 29.39 ± 6.39 |
| | F | 5.55 ± .24 | 5.77 ± .42 | 0.73 | 23.28 ± 2.20 | 25.51 ± 6.59 |
| *Mecp2* | M | 5.69 ± .23 | 4.82 ± .31 | 0.94 | 23.53 ± 1.77 | 15.68 ± 1.81 |
| | F | 5.25 ± .35 | 5.33 ± .37 | 0.91 | 19.31 ± 3.09 | 19.86 ± 3.27 |
| Autism Mutants |  |  |  |  |  |  |
| *Cntnap2* | M | 6.03 ± .37 | 5.56 ± .28 | 0.72 | 28.28 ± 1.95 | 23.95 ± 1.71 |
| | F | 5.65 ± .25 | 5.41 ± .29 | 0.72 | 22.69 ± 2.05 | 19.40 ± 0.97 |
| *Fmr1* | M | 6.03 ± .37 | 6.17 ± .20 | 0.77 | 28.28 ± 1.95 | 29.49 ± 1.37 |
| | F | 5.65 ± .25 | 5.66 ± .17 | 0.47 | 22.69 ± 2.05 | 21.01 ± 1.12 |
| *Shank3* | M | 6.03 ± .37 | 6.00 ± .11 | 0.75 | 28.28 ± 1.95 | 28.24 ± 1.00 |
| | F | 5.65 ± .25 | 5.60 ± .23 | 0.70 | 22.69 ± 2.05 | 21.60 ± 1.70 |
| *Del4Aam* | M | 6.42 ± .36 | 6.18 ± .51 | 0.70 | 33.13 ± 3.21 | 23.22 ± 2.43 |
| | F | 5.50 ± .29 | 5.84 ± .49 | 0.70 | 21.29 ± 1.49 | 18.16 ± 2.32 |

#### 3.4.2 Validation using an ALS model

Mice carrying the SOD1-G93A transgene are a preclinical model of ALS with progressive loss of motor neurons [61, 62]. The SOD1-G93A model has been shown to have changes in gait phenotypes, particularly of hindlimbs [63–69]. The most salient phenotypes are an increase in stance time (duty factor), and decreased stride length in an age-dependent manner. However, several other studies have observed opposite results [63, 64, 67, 68], and some have not seen significant gait effects [70]. These studies did not adjust for body size difference or in some cases for stride speed. We tested SOD1-G93A transgenes and appropriate controls at 64 and 100 days, during time of disease onset [63, 65, 68, 69, 71]. We do not see significant differences in body length or weight (Figure S2C, Table 5), but see changes in stride speed(Figure S2B).

Using model M3, we find small changes in phase of tail tip and nose (Figure 4B). Otherwise, we see significant changes in M1 in stride speed, limb duty factor and stride length (Figure 4A, D). These results argue that the major effect of the SOD1 transgene is on stride speed, which leads to changes in stride time and duty factor. Our results are congruent with reports that gait changes may not be the most sensitive preclinical phenotype in this ALS model, and other phenotypes such as visible clinical signs and motor learning tasks such as rotarod are more sensitive measures [67, 70]. In sum, our results validate the statistical model and may help explain some of the discordant results in the literature.

#### 3.4.3 Validation using a Down syndrome model

Down syndrome, caused by trisomy of all or part of chromosome 21, has complex neurological and neurosensorial phenotypes [72]. Although there are a spectrum of phenotypes such as intellectual disability, seizures, strabismus, nystagmus, and hypoacusis, the more noticeable phenotypes are developmental delays in fine motor skills [73, 74]. These are often described as clumsiness or uncoordinated movements [75, 76]. One of the best studied models, Ts65Dn are trisomic for a region of mouse chromosome 16 that is syntenic to human chromosome 21, recapitulates many of the features of Down syndrome [77, 78]. Ts65Dn mice have been studied for gait phenotypes using traditional inkblot footprint analysis or treadmill methods [79–81]. The inkblot analysis showed mice with shorter and more “erratic” and “irregular” gait, similar to motor coordination deficits seen in patients [80]. Treadmill-based analysis revealed further changes in stride length, frequency, some kinetic parameters, and foot print size [81, 82]. These previous analyses have not studied the whole body posture of these mice.

We analyze Ts65Dn mice along with control mice at approximately 10 and 14 weeks (Table 2) and all three linear mixed models M1-M3 found consistent changes. The Ts65Dn mice are not hyperactive in the open field (Figure 4C), although they have increased stride speed (Figures 4A, C). This indicates that the Ts65Dn mice take quicker steps but travel the same distance as controls. After adjusting stride speed and animal size, step width was increased and step and stride lengths were significantly reduced (Figure 4B). In particular, whole body coordination phenotypes were highly affected in the Ts65Dn mice. The amplitude of tail base and tip, and the phase of tail base, tip, and nose were significantly decreased (Figure 4B). We confirmed this with a phase plot of nose and tail tip (Figure 4E). Surprisingly, we found that there were large differences in phase. The tail tip phase peak is near 30% of the stride cycle in controls and close to 60% in mutants at multiple stride speeds (Figure 4E). Similar changes are seen in the phase plot for the nose. In sum, these results confirm previously reported differences in traditional gait measures, and highlight the utility of our novel open field whole body coordination measures in broadening the assayable phenotypic features in models of human disease. Indeed, the most salient feature of the Ts65Dn gait is the alteration of whole body coordination which previously was reported as a qualitative trait using inkblot analysis [80] and is now quantifiable using our methods.

### 3.5 Characterization of autism spectrum disorder-related mutants

To further validate our approach, we investigated gait in four autism spectrum disorder (ASD) mouse models, in addition to *Mecp2*. In humans, gait and posture defects are often seen in ASD and sometimes gait and motor defects precede classical deficiencies in verbal and social communication and stereotyped behaviors [5, 6]. Recent studies indicate that motor changes are often undiagnosed in ASD cases [83]. It is unclear if these differences have genetic etiologies or are secondary to lack of social interactions that may help children develop learned motor coordination [84]. In mouse models of ASD, gait defects have been poorly characterized, and thus we sought to determine if any gait phenotypes occur in four commonly used ASD genetic models, which we characterized with appropriate controls at 10 weeks (Table 3). Similar to the three models with known gait defects, we tested these mutants and controls in the one hour open field assay and extracted gait and posture metrics (Table 1). We modeled the results using the same approach used for gait mutants (M1 and M3 results are presented in Figure 5, M2 results are in Figure S3).

*Cntnap2* is a member of the neurexin gene family which functions as a cell adhesion molecule [85]. Mutations in *Cntnap2* have been linked to neurological disorders such as ASD, schizophrenia, bipolar disorder, and epilepsy [86]. *Cntnap2* knockout mice have previously been shown to have mild gait effects, with increased stride speed leading to decreased stride duration [87]. These mice are significantly smaller in body length and weight than controls (Table 5, Figure S3C). We used model M2 to compare our results to the previous study and found that *Cntnap2* mice show significant differences in a majority of the gait measures (Figure S3A). In the open field, *Cntnap2* mice were not hyperactive (Figure 5C) but showed a markedly increased stride speed (M1, Figure 5A, C). These results argue that the *Cntnap2* mice do not travel more, but take quicker steps when moving, similar to Ts65Dn mice.

Since *Cntnap2* mice are smaller and have faster stride speeds (Figure 5F), we used results from M3 to determine if gait parameters are altered after adjusting for body size and stride speed (Table 5). We found that *Cntnap2* mice were significantly different from controls for a majority of the traditional gait metrics as well as whole body coordination measures (Figure 5A,B). The *Cntnap2* mice have reduced limb duty factor, step length, step width, and highly reduced stride length (Figure 5B, D). The mice also show altered phase of tail tip, base, and nose, as well as significant but small changes in amplitude of tail tip base and nose. Another salient feature of gait in *Cntnap2* mice is the decrease in inter-animal variance compared to controls, particularly for limb duty factor (Fligner-Killeen test, p *<* 0.01), step length (Fligner-Killeen test, p *<* 0.01), and stride length (Fligner-Killeen test, p *<* 0.02) (Figure 5D). This may indicate a more stereotyped gait in these mutants. Combined, these results imply that *Cntnap2* mice are not hyperactive as measured by total distance traveled in the open field, but are hyperactive at the individual stride level. They take quicker steps with shorter stride and step length, and narrower step width. Next, we asked if there exists a lower-dimensional gait space where the Cntnap2 mutants separate from the controls. We performed principal component analysis (PCA) on z-score transformed gait metrics and embedded the animals in a 2D space for visualization. We found the first PC that explained 40% of the total variance to separate the mutants and controls effectively. We plotted the absolute PC loadings to shed light on contributions of gait and posture metrics to PC1. The loadings revealed that most gait metrics contributed to PC1. We found that the gait metrics allow us to distinguish *Cntnap2* from controls (Figure 5E). This analysis shows that *Cntnap2* mice can be distinguished from controls based on its gait patterns in the open field. We report similar analyses with body length and body length + speed adjusted residuals in Figure S6A,B)

Mutations in *Shank3*, a scaffolding postsynaptic protein, have been found in multiple cases of ASD [88]. Mutations in *Fmr1*, an RNA binding protein that functions as a translational regulator, are associated with Fragile X syndrome, the most commonly inherited form of mental illness in humans [89]. Fragile X syndrome has a broad spectrum of phenotypes that overlaps with ASD features [90]. *Del4Aam* mice contain a deletion of 0.39Mb on mouse chromosome 7 that is syntenic to human chromosome 16p11.2 [91]. Copy number variations (CNVs) of human 16p11.2 have been associated with a variety of ASD features, including intellectual disability, stereotypy, and social and language deficits [92]. *Fmr1* mutant mice travel more in the open field (Figure 5C) and have higher stride speed (Figures 5A, C). When adjusted for stride speed and body length (M3) these mice have slight but significant changes in limb duty factor in model M3. *Shank3* and *Del4Aam* are both hypoactive in the open field compared to controls. *Shank3* mice have a significant decrease in stride speed, whereas *Del4Aam* mice have faster stride speeds (5A, C). All three statistical models show a suggestive or significant decrease in step length in both strains. Using M3, we find that *Shank3* have longer step and stride length, whereas *Del4Aam* have shorter steps and strides. In whole body coordination, *Shank3* mice have a decrease in nose phase and *Del4Aam* has an increase in tail tip phase. These results indicate that, even though both *Shank3* and *Del4Aam* are hypoactive in the open field (Figure 5C), *Shank3* takes slower and longer strides and steps, whereas *Del4Aam* takes faster strides with shorter steps and strides (Figure 5F). Both mutants have some defects in whole body coordination. In sum, we find each of the ASD models to have unique set of gait deficits, with *Cntnap2* having the strongest phenotypes. All have some change in stride speed, although the directionality of change and the variance of the phenotypes differ. These results imply that changes in gait and whole body coordination are general features of ASD.

### 3.6 Strain Survey

After validation of our methods, we sought to understand the range of gait and posture phenotypes in the open field in standard laboratory mouse strains. We surveyed 44 classical inbred laboratory strains, 7 wild derived inbred strains, and 11 F1 hybrid strains (1898 animals, 1,740 hours of video). All animals were isogenic and we surveyed both males and females in a one hour open field assay (Table 4) [42]. We then extracted gait metrics from each video and performed an exploratory analysis of the data on a per-animal level (Figure 6A, S4, S5). We analyzed stride data when animals were traveling at a moderate stride speed (20 to 30 cm/sec) and in a straight direction (angular velocity between -20 to +20 degrees/sec). We could carry out such a selective analysis because of the large amount of data we were able to collect and process in freely moving mice. Since these mice vary considerably in their size [42], we first plotted residuals from M1 that adjusts for body size (Figure 6A, S4, S5).

**Figure 6:**
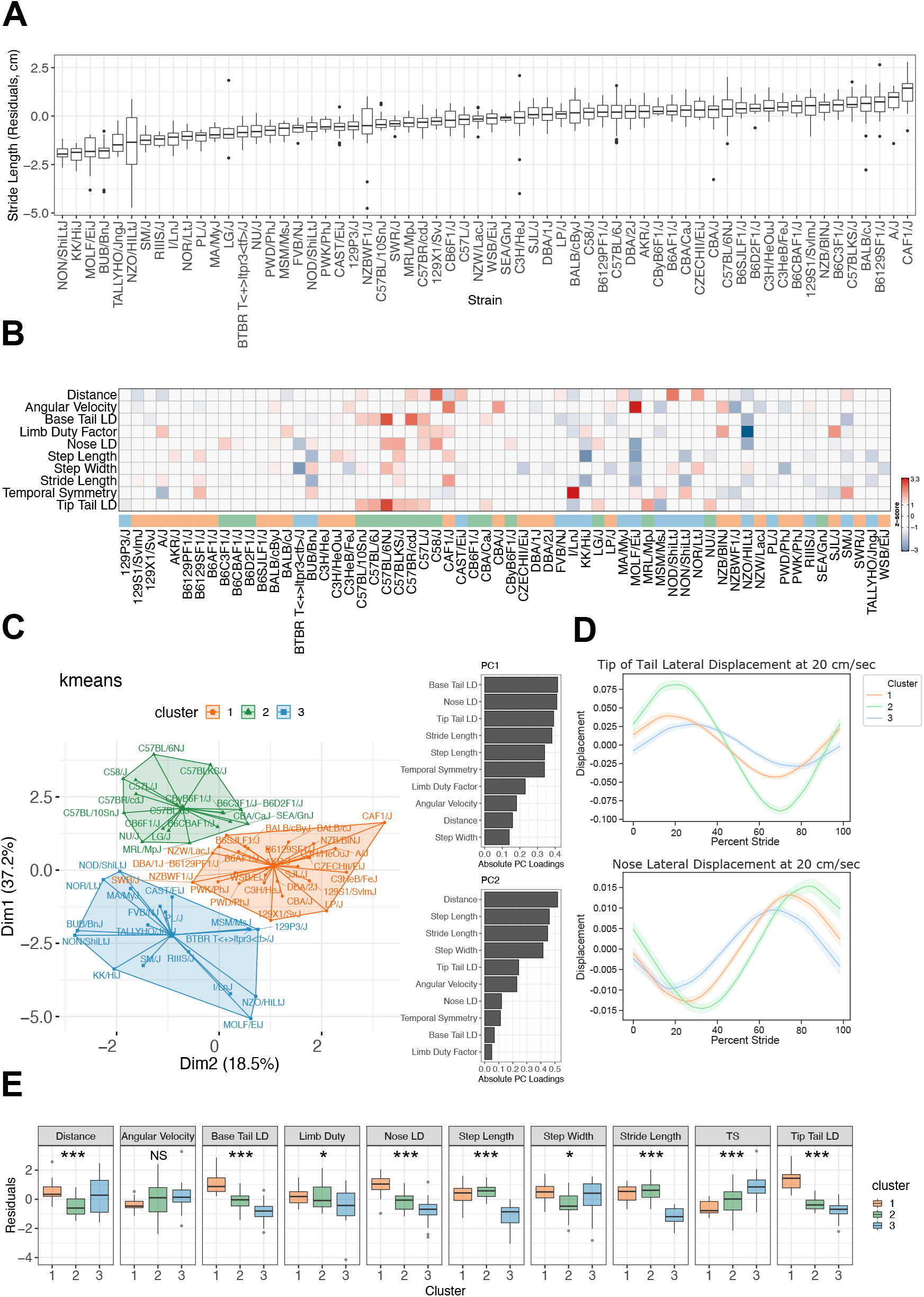
(A) We plot body length adjusted residuals for gait parameter stride length (y-axis) across 62 strains in the strain survey (x-axis). We arrange the boxplots in increasing order of medians from left to right. (B) Residuals are obtained from a linear model with body length and speed as covariates/features for all gait parameters. We transformed the residuals to obtain z-scores and used the scores as input to the next step’s k-means algorithm. The heatmap shows the z-scores (|*z* − score |*>* 1, thresholding is applied for easier visualization) along with color-coded cluster memberships (x-axis). (D) We use a k-means algorithm to determine the cluster memberships. We project the clusters discovered by k-means to a 2D space formed by PC components obtained from the z-scores. See Figure S6 for more information on the choice of the number of clusters and the clusters formed when the gait parameters are adjusted for both body length and body length + stride speed. (D) A consensus view of lateral displacement of nose and tail tip across the clusters. The solid lines represent the mean displacement of stride, while the translucent bands provide a 95% confidence interval for the mean. (E) Post-clustering analysis: We use a linear model (one-way ANOVA) with cluster membership as a categorical covariate/feature to compare gait parameters between strains across the three clusters. The number of *\**s represents the strength of evidence against the null hypothesis of no difference in a gait parameter between strains across 3 clusters. In contrast, NS represents not sufficient evidence to claim a difference in a gait parameter between strains across three clusters.

We sought to determine if we could cluster strains based on their open field gait and posture phenotypes. We took a model-free approach and applied the k-means algorithm to cluster the strains. We did not include the circular features in our analyses as the k-means algorithm requires features that lie in a Euclidean space. We fit a linear model to each linear gait feature with body length and stride speed as covariates and extracted the model residuals. The z-score transformed residuals served as the input features for our analysis (Figure 6B). We initialized the k-means algorithm several times with random points from the data as means. We picked the initialization that gave the smallest total within-cluster sum of squares. We projected the selected k-means output onto the 2D PC subspace to visualize the clustering structure (Figure 6C, D, E). We found three clusters of strains that can be distinguished based on their open field gait behaviors. The number of clusters was chosen based on the gap statistic (Figure S6D). Cluster 1 consisted of primarily classical strains such as A/J, C3H/HeJ, 129S1/SvImJ; cluster 3 consisted of several classical strains and a large number of wild-derived strains such as MOLF/EiJ and CAST/EiJ. Cluster 2 mainly consisted of C57 and related strains, including the reference C57BL/6J. Next, we visualized the clustering structure in a non-linear embedded space using Uniform Manifold Approximation and Projection (UMAP) [93] with two different initializations (Figure S6E). The UMAP dimensions preserved the cluster structure discovered using the k-means algorithm. We further explored the three K-means clusters. PC loading analysis revealed that most of the PC1 is highly correlated with whole body coordination measures while PC2 is correlated with open field distance and traditional gait measures (Figure 6C right). We constructed a consensus stride phase plot of the nose and tail tip for each cluster. Cluster 3 has much higher amplitude, while clusters 1 and 2 have similar amplitude but shifted phase offset (Figure 6D). An examination of the linear gait metrics reveals individual metrics that distinguish the clusters (Figure 6E). For example, cluster 2 has longer stride and step length, while cluster 3 has higher temporal symmetry, while cluster 2 has low lateral displacement of nose, base and tip tail. An overall analysis of individual metrics reveal a significant difference in 9 of 10 measures. For comparison, we plotted the output of the k-means analysis of body length-adjusted gait features (Figure S6C). We found strain SEA/GnJ changed cluster membership from Cluster 2 to Cluster 1 and strains 129X1/SvJ, DBA/2J, LP/J, PWD/PhJ, PWK/PhJ, SWR/J, and WSB/EiJ changed from cluster 1 to cluster 3. In sum, this analysis reveals high levels of heritable variation in gait and whole body posture in the laboratory mouse. A combined analysis using multidimensional clustering of these metrics finds three subtypes of gait in the laboratory mouse. Our results also show that the reference mouse strain, C57BL/6J, is distinct from other common mouse strains and wild derived strains.

We further explored cluster 2 which contains mostly stains from C57 family. We asked whether our movement phenotypes could distinguish among the C57 family. We used two approaches: a supervised dimension reduction approach and a classification approach. For the former, we used linear discriminant analysis (LDA) [26, 94] to quantitatively distinguish between strains C57BL/6J, C57BL/6NJ, C57BLKS/J, C57L/J, C57BR/cdJ, C57BL/10SnJ, and C58/J. C57BL/6J and C57BL/6NJ are considered substrains, while the rest are independent, yet closely related strains [95]. For the second approach, we used a multi-class logistic regression (‘one versus rest’) model to predict the strain membership for each animal from its gait metrics. We adjusted the gait metrics for both body length (Figure S8 A,B) and body length + stride speed (Figure S8 C,D) in our analyses to account for their effect on gait metrics. We found that LDA separated strains when we embedded their adjusted gait metrics in a lower-dimensional 2D space using principal components (Figure S8 A,C). We plotted absolute PC loadings to understand the gait and posture metrics contributions and find that LD1 consists mainly of base tail lateral displacement and LD2 consists of several gait and posture metrics (Figure S8). In the second approach, we summarized the sensitivity of gait analysis for the multi-class classifier to distinguish between strains using a confusion matrix which shows the proportion of correctly classified and misclassified animals in each strain (Figure S8 B,D). Indeed, combined these data indicate that animal movement alone can accurately distinguish genetically similar strains and even substrains.

### 3.7 GWAS

The strain survey demonstrated that the gait features we measure are highly variable, and we thus wanted to understand the heritable components and the genetic architecture of mouse gait in the open field. In human GWAS, both mean and variance of gait traits are highly heritable [96]. We separated the strides of each animal into four different bins according to the speed it was travelling at (10-15, 15-20, 20-25, and 25-30 cm/s) and calculated mean and variance of each trait for each animal in order to conduct a GWAS to identify Quantitative Trait Loci (QTL) in the mouse genome. We used GEMMA [97] to conduct a genome wide association analysis using a linear mixed model, taking into account sex and body length as fixed effects, and population structure as a random effect. Since linear mixed models do not handle circular values, we excluded phase gait data from our analysis. The heritability was estimated by determining the proportion of variance of a phenotype that is explained by the typed genotypes (PVE) (Figure 7A left panel). Heritability of gait measures showed a broad range and the majority of the phenotypes are moderately to highly heritable. The mean phenotypes with lowest heritability are angular velocity and temporal symmetry, indicating that variance in the symmetrical nature of gait or turning behaviors are not due to genetic variance in the laboratory mouse. In contrast, we find that measures of whole body coordination (amplitude measures) and traditional gait measures are moderately to highly heritable. Variance of phenotypes showed moderate heritability, even for traits with low heritability of mean traits (Figure 7A right panel). For instance, mean AngularVelocity phenotypes have low heritability (PVE < 0.1), whereas the variance AngularVelocity phenotypes have moderate heritability (PVE between 0.25 -0.4). These heritability results indicated that the gait and posture traits are appropriate for GWAS of mean and variance traits. Traits with low heritability (PVE < 0.25) were excluded from the GWAS.

**Figure 7:**
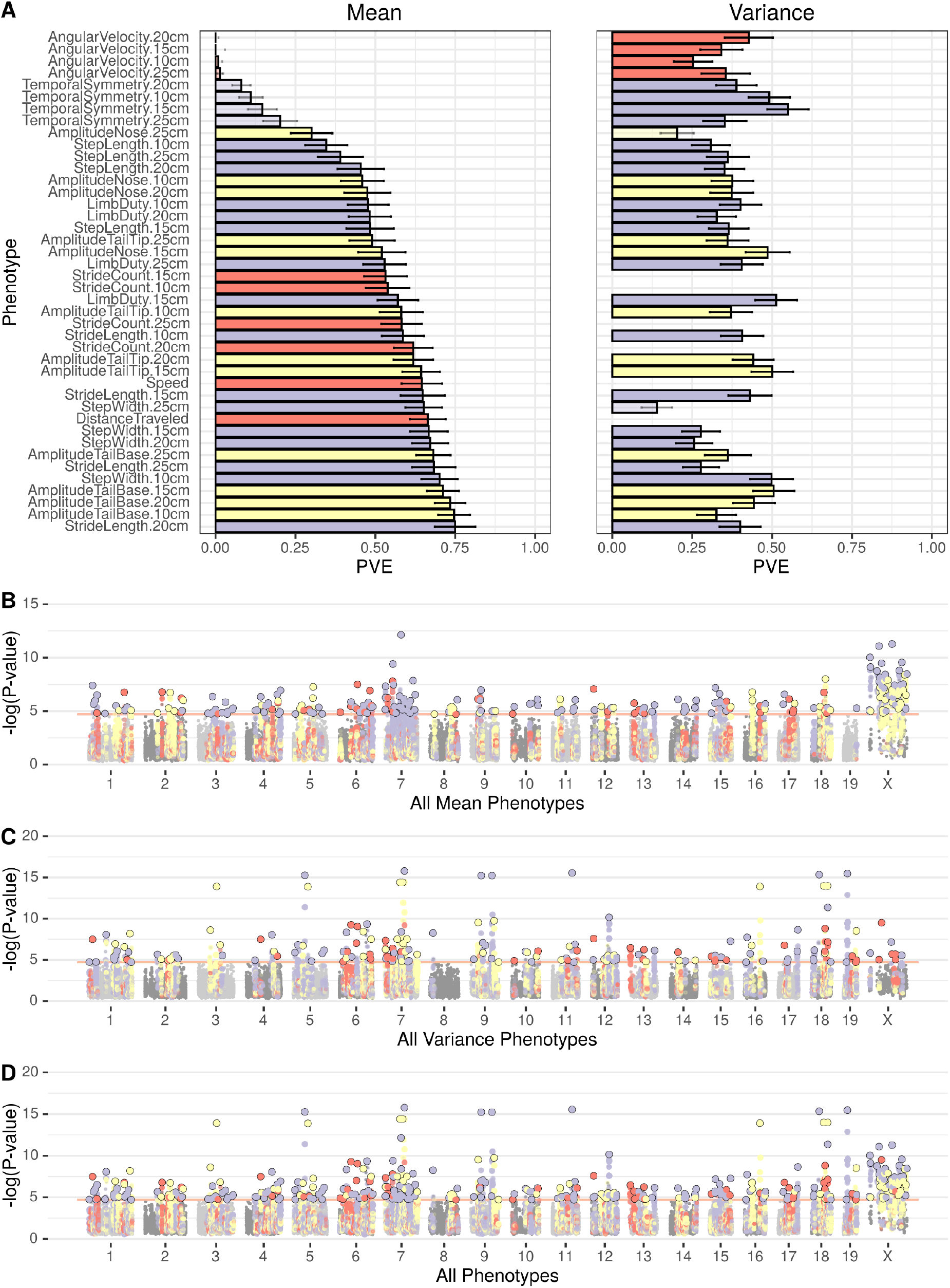
GWAS results for gait phenotypes. (A) Heritability estimates for each phenotype mean (left) and variance (right). Heritability is calculated as PVE (Percent variance explained) (B-D) Manhattan plots of all mean phenotypes (B), variance phenotypes (C), and all of them combined (D); colors correspond to the phenotype with the lowest p-value for the SNP.

For significance threshold, we calculated an empirical p-value correction for the association of a SNP with a phenotype by shuffling the values (total distance traveled in the open field) between the individuals 1000 times. In each permutation, we extracted the lowest p-value to find the threshold that represents a corrected p-value of 0.05 (1.9 *×*10^*−*5^). We took the minimal p-value over all mean phenotypes, variance phenotypes, and both classes combined for each SNP to generate combined Manhattan plots (Figure 7B-D). Each SNP is colored according to the phenotype associated to the SNP with the lowest p-value.

We found 234 QTL for mean traits and 174 QTL for variance traits (Figure 7B-C). The phenotype with the most associated loci was stride length (20-25cm) with 127 loci. Overall, when considering all the phenotypes together, we found 346 significant genomic regions associated with at least one phenotype (Supp. Table 3), indicating only 62 QTL were identified for both a mean phenotype and a variance phenotype. Most phenotypes had limited to no overlap between QTL associated with the mean of the feature and its variance. These data argue that the genetic architecture of mean and variance traits in the mouse are largely independent. These results also begin to outline the genetic landscape of mouse gait and posture in the open field.

We extracted the genes residing in the identified QTL and tested for enriched GO terms, KEGG pathways or Mammalian Phenotypes (MP) associated with them using the software INRICH [98]. Among the most enriched terms are the GO term “ATP-dependent microtubule motor activity, minus-end-directed (MF)” (GO:0008569) with three QTL for stride length containing genes associated with this term out of 20 genes in the genome, and “electron transport chain (BP)” (GO:0022900) enriched in the QTL for step width (Supplementary Figure S9). Since many systems contribute to gait and posture we find over-representation of diverse GO terms. The list of genes found in the QTL interval can be found in supplementary Table S3, genes in highly significant QTL (LOD > 10) were manually annotated for potential impact on gait phenotypes. Of these top 99 genes, most of have been shown to be associated with behavior or neurological, mortality, or growth phenotypes. Only 11 have no reported phenotypes with the rest linked to various developmental and behavioral deficits.

## 4 Discussion

Gait and posture are an important indicator of health, and are perturbed in many neurological, neuromuscular, and neuropsychiatric diseases. The goal of this project was to develop a simple and reliable automated system that is capable of performing pose estimation on mice and to extract key gait and posture metrics from pose. We present a solution that allows researchers to adapt a traditional video imaging system used for open field analysis to extract gait metrics. Our approach has some clear advantages as well as limitations. We are able to process a large amount of data with low effort and low cost since the only data that needs to be captured is top-down gray scale video of a mouse in an open field, and all pose estimation and gait metric extraction is fully automated after that. Top down videos have routinely been used in behavioral neurogenetics and this method could be applied to archival video data. Indeed, we analyzed gait in a strain survey data set that we partially analyzed for tracking and grooming behaviors [42, 99]. Because our method does not require expensive specialized equipment, we can also allow the mouse time to acclimate to the open field and collect data over long periods of time, potentially over days with the addition of bedding, food, and water. Additionally our method allows the animal to move of its own volition (unforced behavior) in a familiar environment, a more ethologically relevant assay [32]. One limitation of our approach is that we cannot measure kinetic properties of gait because we are limiting ourselves to video [27]. We also limit ourselves to a 2D representation of pose because of our monocular recording configuration. A 3D representation of pose would allow for the extraction of height metrics from all keypoints and would likely provide richer gait phenotypes [41]. The decision to use top-down video also means that forepaw keypoints are often occluded by the mouse’s body. The pose estimation network is robust to some amount of occlusion as is the case with the hind paws but the forepaws, which are almost always occluded during locomotion, have pose estimates which are too inaccurate and so have been excluded from our analysis. Regardless, in all genetic models that we tested, hind paw data is sufficient to detect robust differences in gait and body posture. In addition, we analyze videos at 30hz (frames per second) which is standard for video streams. Certain behaviors that occur at high speed such as escape or gallop may be difficult to determine. Other more kinematic approaches that view the animal from multiple angles and capture data at high frame rates may be more appropriate for certain applications [25, 26]. Thus, our methods are not a replacement for kinematic gait analysis that is carried out by certain specialized labs. These labs need higher resolution approaches with enhanced video and kinetic analysis. Instead, our method offers a high-throughput assessment of gait in a commonly used behavior apparatus, the open field. We hope that our methods will allow these behavior labs to easily access gait and posture for additional biological insight. In addition, the ability to analyze large amounts of data in free moving animals, proves to be highly sensitive, even with very strict heuristic rules around what we consider to be a gait. Future iterations of our method could incorporate data from multiple camera angles and with higher frame rate.

The gait measures we extract are commonly quantified in experiments (e.g. step width and stride length), but measures of whole body coordination such as lateral displacement and phase of tail are typically not measured in rodent gait experiments (phase and amplitude of keypoints during stride). Gait and whole body posture is frequently measured in humans as an endophenotype of psychiatric illness [2, 5, 7, 8]. Our results in mice indicate that gait and whole body coordination measures are highly heritable and perturbed in disease models. Specifically, we test neurodegenerative (*Sod1*), neurodevelopmental (Down syndrome, *Mecp2*) and ASD models (*Cntnap2, Shank3, FMR1, Del4Am*) and find altered gait features in all of these mutants. Others have also found similar results with neurodegerative models [25]. Of note are the data for Down syndrome. In humans, miscoordination and clumsiness are prominent features of Down syndrome. In mouse models, this miscoordination was previously characterized in ink blot gait assays as a disorganized hind foot print. Here, our analysis revealed perturbed whole body coordination differences between control and Tn65Dn mice. Our approach thus enables quantitation of a previously qualitative trait.

Our analysis of a large number of mouse strains for gait and posture finds three distinct classes of overall movement. We find that the reference C57BL/6J and related strains belong to a distinct cluster separate from other common laboratory as well as wild-derived strains. The main difference is seen in the high amplitude of tail and nose movement of the C57BL/6 and related strains. This may be important when analyzing gait and posture in differing genetic backgrounds. The GWAS revealed 346 QTL (including 62 pleiotropic associations) for gait and posture in the open field for both mean and variance phenotypes. We found that the mean and variance of traits are regulated by distinct genetic loci. Indeed, we find most variance phenotypes show moderate heritability, even for mean traits with low heritability. Human GWAS have been conducted for gait and posture, albeit with under powered samples, which has lead to good estimates of heritability but only a few significantly associated loci [96]. Enrichment analysis showed loose set of GO terminologies that are enriched, indicating a wide array of biological functions that regulates gait and posture. Our results in the mouse argue that a well-powered study in humans may identify hundreds of genetic factors that regulates gait and posture.

## 5 Methods

### 5.1 Animals and behavioral testing

All behavioral tests were performed in accordance with approved protocols from The Jackson Laboratory Institutional Animal Care and Use Committee guidelines. All animals were obtained from JAX repository or bred in a room adjacent to the behavioral testing room Table 5 and 4. All behavioral protocols have been previously published [42, 100]. The strain survey data was published before and reanalyzed for gait behavior [42, 99]. We excluded animals that had too few strides which disproportionately affected low activity strains, leading to the final animal numbers indicted in Table 4. All gait mutants and ASD models were generated in JAX repository and genotyped prior to shipment to Kumar Lab for testing. Animals were acclimated for at least a week prior to any testing. Prior to open field testing, animals were moved to the behavior room and allowed to acclimate for 30-60 minutes. White noise was used for testing to balance the noise between holding and testing rooms.

### 5.2 Data Availability

All code and data will be available on the www.Kumarlab.org website and Github account (https://github.com/KumarLabJax/) upon publication. Behavioral metrics extracted in this manuscript will be deposited on the Mouse Phenome Database (www.phenome.jax.org under dataset Kumar4).

### 5.3 Training Data

Labeled data consists of 8,910 480×480 grayscale frames containing a single mouse in the open field along with the twelve manually labeled pose keypoints per frame. We selected from a diverse set of mouse strain with different appearance accounting for variation in coat color, body size and obesity. Figure 2C shows a representative frame generated by our open field apparatus. The frames were generated from the same open field apparatus as was used to generate experimental data previously [42]. Pose keypoint annotations were performed by several Kumar lab members. Frame images and keypoint annotations were stored together using an HDF5 format which was used for neural network training. Frame annotations were split into a training dataset (7,910 frames) and a validation dataset (1,000 frames) for training.

### 5.4 Neural Network Training

We train our network over 600 epochs and perform validation at the end of every epoch. The training loss curves (figure 2C) show a fast convergence of the training loss without an overfitting of the validation loss. We used transfer learning [101, 102] on our network in order to minimize the labeling requirements and improve the generality of our model. We started with the imagenet model provided by the authors of the HRNet paper (hrnet_w32-36af842e.pth) and froze the weights up to the second stage during training. In order to further improve the generality of our network we employed several data augmentation techniques during training including: rotation, flipping, scaling, brightness, contrast and occlusion. We use the ADAM optimizer to train our network. The learning rate is initially set to 5 *×* 10^*−*4^, then reduced to 5 *×* 10^*−*5^ at the 400^th^ epoch and 5 *×* 10^*−*6^ at the 500^th^ epoch.

### 5.5 Gait Extraction

Here we describe our method of extracting gait structure from pose in further detail. The processes of detecting strides begins with first determining intervals of time where the mouse is moving at sufficient speed for strides to take place. We will call these tracks. To determine track intervals we threshold for base of tail speed greater than or equal to 5 cm/sec. Through observation we determined that the base of tail point is highly stable and a good surrogate for overall mouse speed.

The next step is to identify individual steps in both the left hind paw and right hind paw. Initially these steps are determined for each paw respectively without consideration for the other paw. Later in the process we will pair up left and right steps into strides. The process for step detection relies on oscillations in speed as can be seen in (Figure 2D). We calculate individual paw speed and then apply a peak detection algorithm [103] to identify local maxima in speed. After finding all of the local maxima we use the surrounding local minima to define a step interval with a toe-off event followed by a foot strike event on either side of the step. We then filter out any steps whose peak speed does not exceed 15 cm/sec or the overall animal speed (whichever is greater).

Once we have each set of valid steps from left and right hind paws we need to group pairs of steps together to find strides. We use left hind paw steps to delimit strides. Stride intervals end when the left hind paw step ends and begin at the frame just after the previous stride. There is an additional constraint that stride intervals are not allowed to extend before or after the containing track. After we have defined our stride intervals using left hind paw steps we need to associate right hind paw steps with the stride. If we find a right hind paw step that completes within the given stride interval we then associate that step with the stride. If we cannot find a right hind step that meets this condition the stride is discarded.

Now that we have all of our stride definitions with associated steps we apply additional filtering to improve the quality and consistency of strides. For our application of statistical and genomic analysis of stride metrics we decided to take a fairly aggressive approach at removing strides that have potential to degrade quality or introduce inconsistency. All strides at the start and end of a track are removed. This is done to improve the consistency of gait metrics and avoid introducing variance due to starting and stopping behavior. We also discard strides if the keypoint confidence for Nose, Neck Base, Spine Center, Tail Base, Hind Paw Left, Hind Paw Right, Tail Middle or Tail Tip (Figure 2 G) falls below the threshold of 0.3 for any frame in the stride in order to avoid using low quality strides.

### 5.6 Corner Detection

In order to perform a comparative analysis of gait metrics between center and periphery of the open field arena (Figure S7) we need to know pixel locations for each of the four corners. Rather than use fixed values which would have been affected by differences in camera placement, we developed a simple corner detector using the same neural network architecture that we developed for pose estimation. The only difference in the network structure is the output; the pose estimation network outputs twelve heatmaps (one per keypoint) whereas the corner detection network just outputs a single heatmap for detecting corner positions. We trained our corner detector using 415 annotated images. We tested accuracy on the 27 validation images which use the same 480×480 resolution that we use for the video in this paper. The Mean Absolute Error (MAE) averaged over the 108 corners from these 27 validation images was 1.63 ±0.51 pixels (0.20 ±0.06 cm). In order to generate corner positions for each of our videos we choose a frame from seven different time points and estimate corner locations at each of these frames. For each corner we then use the median X and Y values from these seven measurements as the final value in order to reduce the impact of inaccurate locations which can result from occlusion or other image noise.

### 5.7 Statistical Analysis

Each gait and autism mutant were analyzed separately one at a time. Having fixed a mutant line to analyze, we used dummy variable encoding to encode Genotype, a categorical covariate with two levels -Control (0) and Mutant (1), with the corresponding littermate WTs (or another control strain) serving as the reference level. The numeric covariates, Body length (M1,M2,M3), speed (M2, M3), were normalized using the z-score transformation. We did not include Sex as a covariate in the model; we found it correlated with body length. As described earlier, we treated all gait metrics as response variables except in M2 and M3, where we treated stride speed as a covariate. For the linear gait metrics, we considered the following LMM model for repeated measurements:

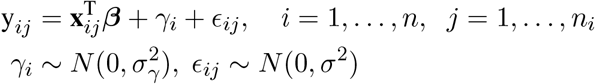

where *n* is the total number of animals; y_*ij*_ is the *j*^th^ repeat measurement on the *i*^th^ animal, *n*_*i*_ denotes the number of repeat measurements on animal *i*; x_*ij*_ is a *p ×* 1 vector of covariates such as body length, stride speed, genotype, age; *β* is a *p ×* 1 vector of unknown fixed population-level effects; *γ*_*i*_ is a random intercept which describes subject-specific deviation from the population mean effect; and *ϵ*_*ij*_ is the error term that describes the intrasubject variation of the *i*^th^ subject that is assumed to be independent of the random effect. We used Type II ANOVA F-test via Satterthwaite’s degrees of freedom method to test the null hypothesis of no Genotype-based effect and obtain p-values. We fit our LMM models using the lme4 package in R [104].

We modeled the circular gait metrics (phase variables) in Table 1 as a function of linear variables using a circular-linear regression model. Analyzing circular data is not straightforward and statistical models developed for linear data do not apply to circular data [105]. The circular response variables are assumed to have been drawn from a von-Mises distribution with unknown mean direction *µ* and concentration parameter *κ*. The mean direction parameter is related to the variables **X** through the equation

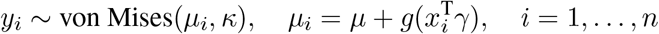

where *y*_*i*_ is the mean circular metric for animal *i, g*(*u*) = 2 tan^*−*1^(*u*) is a link function such that for *−∞< u < ∞,−π < g*(*u*) *< π*. The parameters *µ, γ*_1_, …, *γ*_*k*_ and *κ* are estimated via maximum likelihood [54]. We computed the p-value of no Genotype-based effect in circular phenotypes using a t-test which is based on asymptotic normality of the maximum likelihood estimators of the model parameters. The model is fitted using the circular package in R. [106]

We used the q-value (FDR adjusted p-value) to adjust for multiple testing across all gait metrics and control the false positive discovery rate. We reported LOD scores, defined as -log(q-value), in the heatmaps (Figures 4A,B, 5A,B, S2A, and S3A), and the q-values in Tables S3, S4.

The gait features for each animal were adjusted for body length (resp. body length + stride speed) by extracting residuals from a linear model with body length (resp. body length and stride speed) covariate/s and the gait metrics as response variables. The residuals were then averaged over animals for each strain to form a representative observation for the strain. The input to the k-means algorithm consisted of a matrix of 62 z-score transformed observations, each corresponding to a strain, on ten gait metrics. We initialized the k-means algorithm several times with random points from the data as means. We picked the initialization that gave the smallest total within-cluster sum of squares. We projected the selected k-means output onto the 2D PC (principal components) subspace to visualize the clustering structure. We performed PCA by applying singular value decomposition (SVD) of the input observations matrix. We obtained the contributions of different gait features to each PC component using the PC loadings, i.e., the corresponding eigenvectors obtained from PCA. The number of clusters was chosen based on the gap statistic [107]. To see the effect of non-linear embedding, we visualized the clustering structure in a non-linear embedded space using UMAP [93] with two different initializations (SPCA -scaled PCA, Laplace -Laplacian Eigenmap).

### 5.8 GWAS

The R package *mousegwas*, previously described in [99], was used to compute genome wide associations. Briefly, the classical mouse strains were used (excluding wild-derived strains), ten individuals from each strain and sex combination were randomly selected and GEMMA was used with its LMM method. The MDA genotypes were obtained from (https://phenome.jax.org/genotypes). The GWAS can be reproduced using the command:

~~~
export G=https://raw.githubusercontent.com/TheJacksonLaboratory/mousegwas/mastenextflowrunTheJacksonLaboratory/mousegwas-rgait \
--yaml $G/example/gait_nowild.yaml \
--shufyaml $G/example/gait_shuffle.yaml \
--input $G/example/gait_paper_strain_survey_2019_11_12.csv \
--outdir gait_output --clusters 1 \
--addpostp “--colorgroup --meanvariance --set3 --minherit 0.25 --loddrop 1.5” \
--addheatmap “--meanvariance -p 0.1” -profile slurm,singularity
~~~

## 6 Acknowledgements

We thank members of the Kumar Lab for helpful advice and Taneli Helenius for editing. We thank Massimo Daul for preliminary work on corner detection of our open fields. We thank Dr. Vivek Philip for advice on GWAS analysis. We thank JAX Information Technology team members Edwardo Zaborowski, Shane Sanders, Rich Brey, David McKenzie, and Jason Macklin for infrastructure support. This work was funded by The Jackson Laboratory Directors Innovation Fund, National Institute of Health DA041668 (NIDA), DA048634 (NIDA), and Brain and Behavioral Foundation Young Investigator Award (V.K.). All code and training data will be available at Kumarlab.org and Kumar Lab Github upon publication.

## 8 Supplementary data not in pdf document

Video S1 -Examples of pose estimation. Pose estimation on visually diverse mouse strains using the architecture shown in Figure 1.

Video S2 -Gait extraction from pose estimation. The top panel shows a segment of video with a gait overlay similar to what is described in Figure 2C. The bottom panel contains two plots that update with the video: an angular velocity plot similar to Figure 2F and a hind paw speed plot as described in Figure 2D. Green bouts are considered strides, left and right paws are orange and blue respectively.

Video S3 -Aggregated stride cycles for the *Ts65Dn Trisomic* mouse model: individual stride cycle keypoints overlaid and rendered as point clouds (left), and 2D density plots of stride cycle keypoints (right) of gait in a stride cycle. Data are combined across all Ts65Dn Trisomic mice within the 20 to 25 cm/sec speed bin and -20 to 20 deg/sec angular velocity bin. Each dot represents keypoint from one stride on left. The colors denote the various keypoints.

Video S4 -NOR/LtJ gait video clip (see Figure 3).

Video S5 -C57BL/6J gait video clip (see Figure 3).

Video S6 -Overlay of NOR/LtJ and C57BL/6J stride cycle showing antiphasic whole body posture (see Figure 3). In order to generate this animation data are combined across all NOR/LtJ and across all C57BL/6J within the strain survey for the 20 to 25 cm/sec speed bin and -20 to 20 deg/sec angular velocity bin. Each point represents the median X and Y position for each strain at the frame’s time point.

Supplementary table 3 -QTL table from GWAS analysis and Analysis of top gene candidates in peak that have LOD scores over 10.

## 9 Supplement

**Figure S1:**
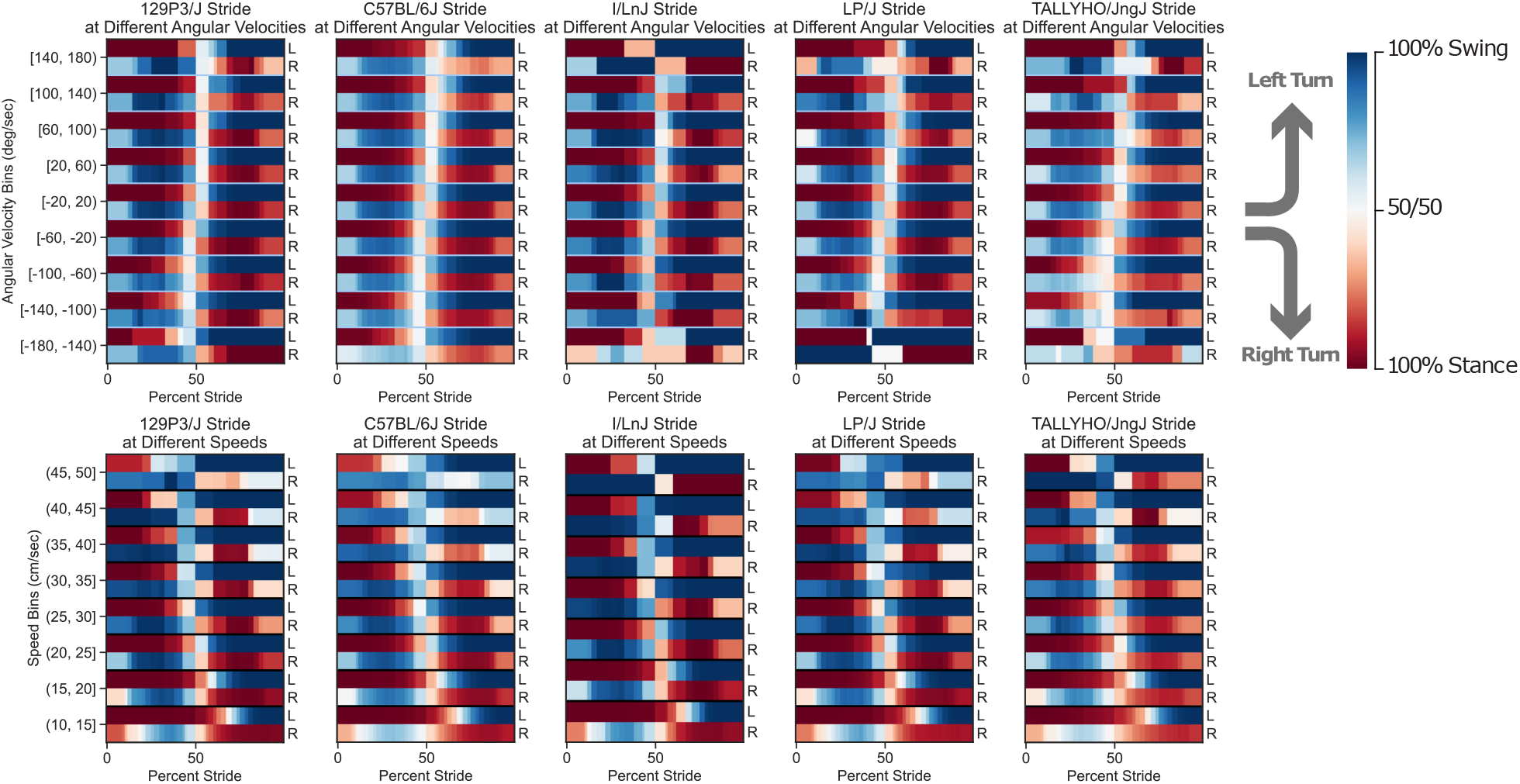
Hildebrand plots for hind paws over five diverse strains organized in columns. Each of these is aggregated over multiple strides from multiple individuals. The top row is separated into different angular velocity bins along the x axis showing how the stride pattern changes with angular velocity and the x axis for the bottom row is separated into stride speed bins showing how the stride pattern changes with speed.

**Table S1:** We show errors in pixels and centimeters given for all keypoints grouped together and broken out for specific keypoint types. We use 1000 of our validation images to calculate Mean Absolute Error (MAE) with a Standard Error of Mean (SEM). We also calculate MEA and SEM for 200 dark colored mice and 200 white mice to demonstrate that our pose estimation network is robust to visual difference.

| Keypoint | MAE ± SEM (Pixels) | | | MAE ± SEM (cm) | | |
| --- | --- | --- | --- | --- | --- | --- |
|  | All 1000 | Dark | White | All 1000 | Dark | White |
| All Keypoints | 3.27 ± 0.05 | 3.34 ± 0.06 | 3.01 ± 0.05 | 0.41 ± 0.01 | 0.41 ± 0.01 | 0.37 ± 0.01 |
| Nose | 2.53 ± 0.10 | 2.50 ± 0.16 | 2.23 ± 0.11 | 0.31 ± 0.01 | 0.31 ± 0.02 | 0.28 ± 0.01 |
| Ear Left | 2.41 ± 0.07 | 2.39 ± 0.12 | 2.12 ± 0.10 | 0.30 ± 0.01 | 0.30 ± 0.02 | 0.26 ± 0.01 |
| Ear Right | 2.18 ± 0.07 | 1.90 ± 0.10 | 2.08 ± 0.11 | 0.27 ± 0.01 | 0.24 ± 0.01 | 0.26 ± 0.01 |
| Neck Base | 2.39 ± 0.05 | 2.21 ± 0.10 | 2.24 ± 0.09 | 0.30 ± 0.01 | 0.27 ± 0.01 | 0.28 ± 0.01 |
| Forepaw Left | 4.14 ± 0.09 | 4.63 ± 0.19 | 4.44 ± 0.19 | 0.51 ± 0.01 | 0.57 ± 0.02 | 0.55 ± 0.02 |
| Forepaw Right | 4.08 ± 0.09 | 4.79 ± 0.20 | 4.05 ± 0.17 | 0.50 ± 0.01 | 0.59 ± 0.02 | 0.50 ± 0.02 |
| Spine Center | 3.66 ± 0.07 | 4.41 ± 0.16 | 3.17 ± 0.14 | 0.45 ± 0.01 | 0.55 ± 0.02 | 0.39 ± 0.02 |
| Hind Paw Left | 5.08 ± 0.15 | 6.38 ± 0.36 | 3.68 ± 0.25 | 0.63 ± 0.02 | 0.79 ± 0.04 | 0.46 ± 0.03 |
| Hind Paw Right | 4.07 ± 0.13 | 3.79 ± 0.30 | 4.81 ± 0.28 | 0.50 ± 0.02 | 0.47 ± 0.04 | 0.60 ± 0.03 |
| Tail Base | 2.70 ± 0.09 | 2.90 ± 0.23 | 2.73 ± 0.16 | 0.33 ± 0.01 | 0.36 ± 0.03 | 0.34 ± 0.02 |
| Tail Middle | 3.24 ± 0.12 | 2.70 ± 0.14 | 2.62 ± 0.14 | 0.40 ± 0.01 | 0.33 ± 0.02 | 0.32 ± 0.02 |
| Tail Tip | 2.77 ± 0.51 | 1.53 ± 0.14 | 1.91 ± 0.14 | 0.34 ± 0.06 | 0.19 ± 0.02 | 0.24 ± 0.02 |

**Table S2:**
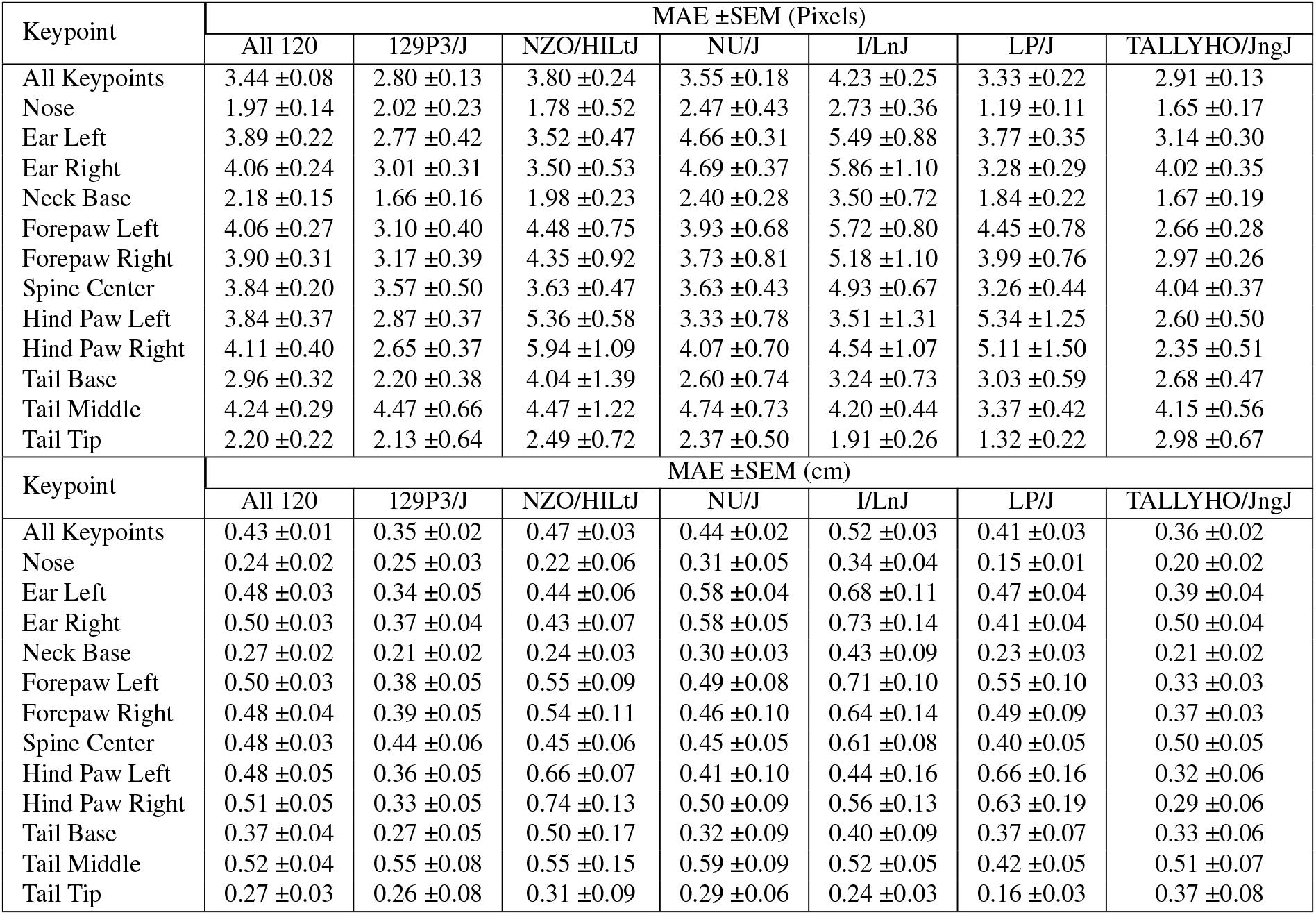
We show errors in pixels and centimeters given for all keypoints grouped together and broken out for specific keypoint types. We use 120 validation images (20 from each of the six strains shown) to calculate Mean Absolute Error (MAE) with a Standard Error of Mean (SEM). The mice chosen vary significantly in appearance in order to demonstrate the networks robustness to diversity. We include off-white (129P3/J), black obese (NZO/HILtJ), nude (NU/J), piebald (I/LnJ), agouti (LP/J) and moderately obese albino (TALLYHO/JngJ) mice. These are the same strains for which we render pose estimation in our supplementary video: VideoS1_diverse-mouse-pose.mp4

| Keypoint | MAE ±SEM (Pixels) | | | | | | |
| --- | --- | --- | --- | --- | --- | --- | --- |
|  | All 120 | 129P3/J | NZO/HILtJ | NU/J | I/LnJ | LP/J | TALLYHO/JngJ |
| All Keypoints | 3.44 ±0.08 | 2.80 ±0.13 | 3.80 ±0.24 | 3.55 ±0.18 | 4.23 ±0.25 | 3.33 ±0.22 | 2.91 ±0.13 |
| Nose | 1.97 ±0.14 | 2.02 ±0.23 | 1.78 ±0.52 | 2.47 ±0.43 | 2.73 ±0.36 | 1.19 ±0.11 | 1.65 ±0.17 |
| Ear Left | 3.89 ±0.22 | 2.77 ±0.42 | 3.52 ±0.47 | 4.66 ±0.31 | 5.49 ±0.88 | 3.77 ±0.35 | 3.14 ±0.30 |
| Ear Right | 4.06 ±0.24 | 3.01 ±0.31 | 3.50 ±0.53 | 4.69 ±0.37 | 5.86 ±1.10 | 3.28 ±0.29 | 4.02 ±0.35 |
| Neck Base | 2.18 ±0.15 | 1.66 ±0.16 | 1.98 ±0.23 | 2.40 ±0.28 | 3.50 ±0.72 | 1.84 ±0.22 | 1.67 ±0.19 |
| Forepaw Left | 4.06 ±0.27 | 3.10 ±0.40 | 4.48 ±0.75 | 3.93 ±0.68 | 5.72 ±0.80 | 4.45 ±0.78 | 2.66 ±0.28 |
| Forepaw Right | 3.90 ±0.31 | 3.17 ±0.39 | 4.35 ±0.92 | 3.73 ±0.81 | 5.18 ±1.10 | 3.99 ±0.76 | 2.97 ±0.26 |
| Spine Center | 3.84 ±0.20 | 3.57 ±0.50 | 3.63 ±0.47 | 3.63 ±0.43 | 4.93 ±0.67 | 3.26 ±0.44 | 4.04 ±0.37 |
| Hind Paw Left | 3.84 ±0.37 | 2.87 ±0.37 | 5.36 ±0.58 | 3.33 ±0.78 | 3.51 ±1.31 | 5.34 ±1.25 | 2.60 ±0.50 |
| Hind Paw Right | 4.11 ±0.40 | 2.65 ±0.37 | 5.94 ±1.09 | 4.07 ±0.70 | 4.54 ±1.07 | 5.11 ±1.50 | 2.35 ±0.51 |
| Tail Base | 2.96 ±0.32 | 2.20 ±0.38 | 4.04 ±1.39 | 2.60 ±0.74 | 3.24 ±0.73 | 3.03 ±0.59 | 2.68 ±0.47 |
| Tail Middle | 4.24 ±0.29 | 4.47 ±0.66 | 4.47 ±1.22 | 4.74 ±0.73 | 4.20 ±0.44 | 3.37 ±0.42 | 4.15 ±0.56 |
| Tail Tip | 2.20 ±0.22 | 2.13 ±0.64 | 2.49 ±0.72 | 2.37 ±0.50 | 1.91 ±0.26 | 1.32 ±0.22 | 2.98 ±0.67 |
| Keypoint | MAE ±SEM (cm) | | | | | | |
|  | All 120 | 129P3/J | NZO/HILtJ | NU/J | I/LnJ | LP/J | TALLYHO/JngJ |
| All Keypoints | 0.43 ±0.01 | 0.35 ±0.02 | 0.47 ±0.03 | 0.44 ±0.02 | 0.52 ±0.03 | 0.41 ±0.03 | 0.36 ±0.02 |
| Nose | 0.24 ±0.02 | 0.25 ±0.03 | 0.22 ±0.06 | 0.31 ±0.05 | 0.34 ±0.04 | 0.15 ±0.01 | 0.20 ±0.02 |
| Ear Left | 0.48 ±0.03 | 0.34 ±0.05 | 0.44 ±0.06 | 0.58 ±0.04 | 0.68 ±0.11 | 0.47 ±0.04 | 0.39 ±0.04 |
| Ear Right | 0.50 ±0.03 | 0.37 ±0.04 | 0.43 ±0.07 | 0.58 ±0.05 | 0.73 ±0.14 | 0.41 ±0.04 | 0.50 ±0.04 |
| Neck Base | 0.27 ±0.02 | 0.21 ±0.02 | 0.24 ±0.03 | 0.30 ±0.03 | 0.43 ±0.09 | 0.23 ±0.03 | 0.21 ±0.02 |
| Forepaw Left | 0.50 ±0.03 | 0.38 ±0.05 | 0.55 ±0.09 | 0.49 ±0.08 | 0.71 ±0.10 | 0.55 ±0.10 | 0.33 ±0.03 |
| Forepaw Right | 0.48 ±0.04 | 0.39 ±0.05 | 0.54 ±0.11 | 0.46 ±0.10 | 0.64 ±0.14 | 0.49 ±0.09 | 0.37 ±0.03 |
| Spine Center | 0.48 ±0.03 | 0.44 ±0.06 | 0.45 ±0.06 | 0.45 ±0.05 | 0.61 ±0.08 | 0.40 ±0.05 | 0.50 ±0.05 |
| Hind Paw Left | 0.48 ±0.05 | 0.36 ±0.05 | 0.66 ±0.07 | 0.41 ±0.10 | 0.44 ±0.16 | 0.66 ±0.16 | 0.32 ±0.06 |
| Hind Paw Right | 0.51 ±0.05 | 0.33 ±0.05 | 0.74 ±0.13 | 0.50 ±0.09 | 0.56 ±0.13 | 0.63 ±0.19 | 0.29 ±0.06 |
| Tail Base | 0.37 ±0.04 | 0.27 ±0.05 | 0.50 ±0.17 | 0.32 ±0.09 | 0.40 ±0.09 | 0.37 ±0.07 | 0.33 ±0.06 |
| Tail Middle | 0.52 ±0.04 | 0.55 ±0.08 | 0.55 ±0.15 | 0.59 ±0.09 | 0.52 ±0.05 | 0.42 ±0.05 | 0.51 ±0.07 |
| Tail Tip | 0.27 ±0.03 | 0.26 ±0.08 | 0.31 ±0.09 | 0.29 ±0.06 | 0.24 ±0.03 | 0.16 ±0.03 | 0.37 ±0.08 |

**Figure S2:**
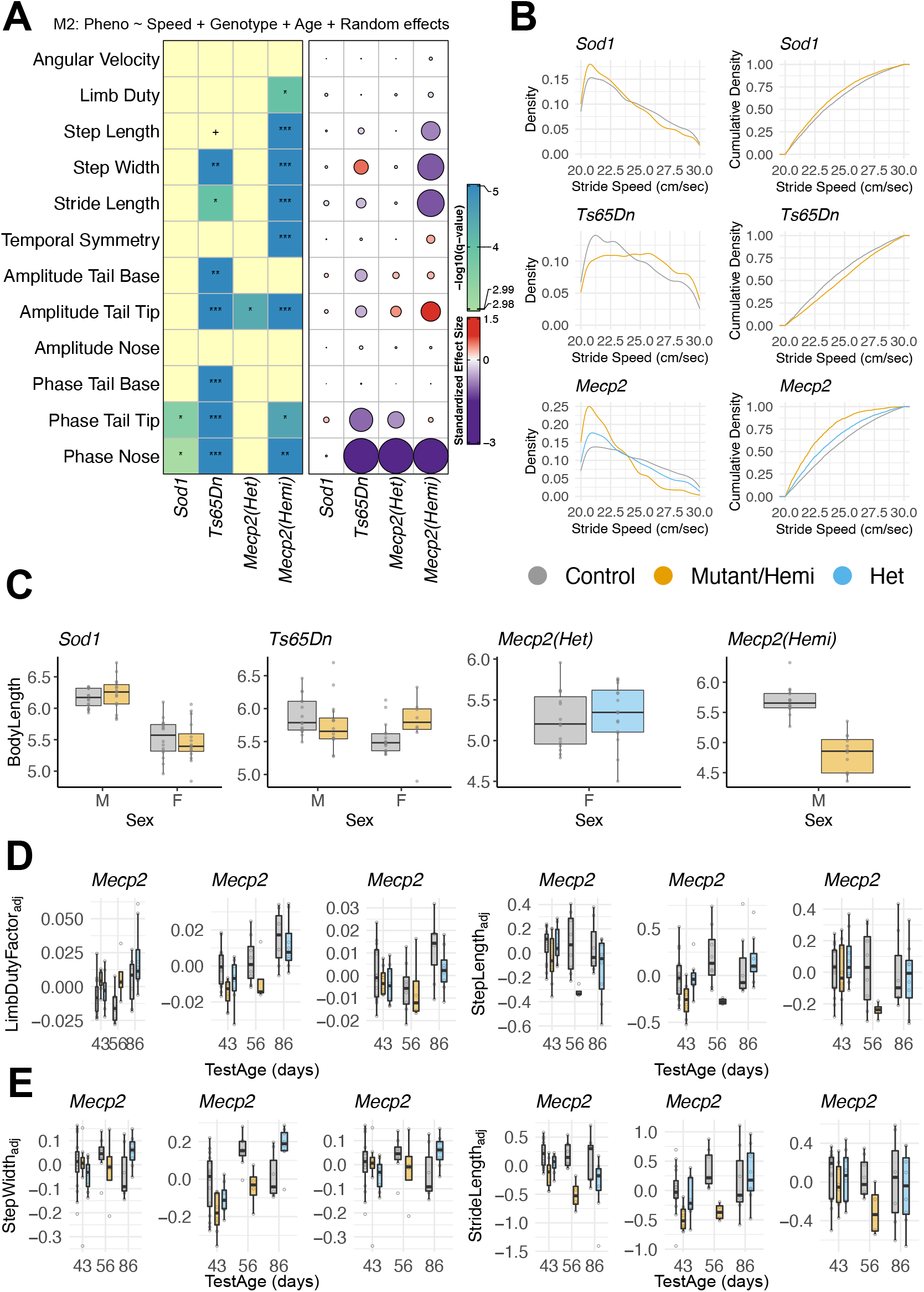
Supplemental analysis of gait mutants described in Figure 4. (A) Heat map summarizing the effect sizes and q-values obtained from model M3: Phenotype *∼* Genotype + TestAge + Speed + BodyLength + (1 | MouseID/TestAge). (B) Kernel density (left) and cumulative density (right) curves of stride speed across all strains. (C) A plot showing positive association between body length and sex across different gait mutant strains. (D) Body length (M1), stride speed (M2), body length and stride speed (M3) adjusted residuals for limb duty factor and step length for Mecp2 gait mutant. (E) Body length (M1), stride speed (M2), body length and stride speed (M3) adjusted residuals for step width and stride length for Mecp2 gait mutant.

**Figure S3:**
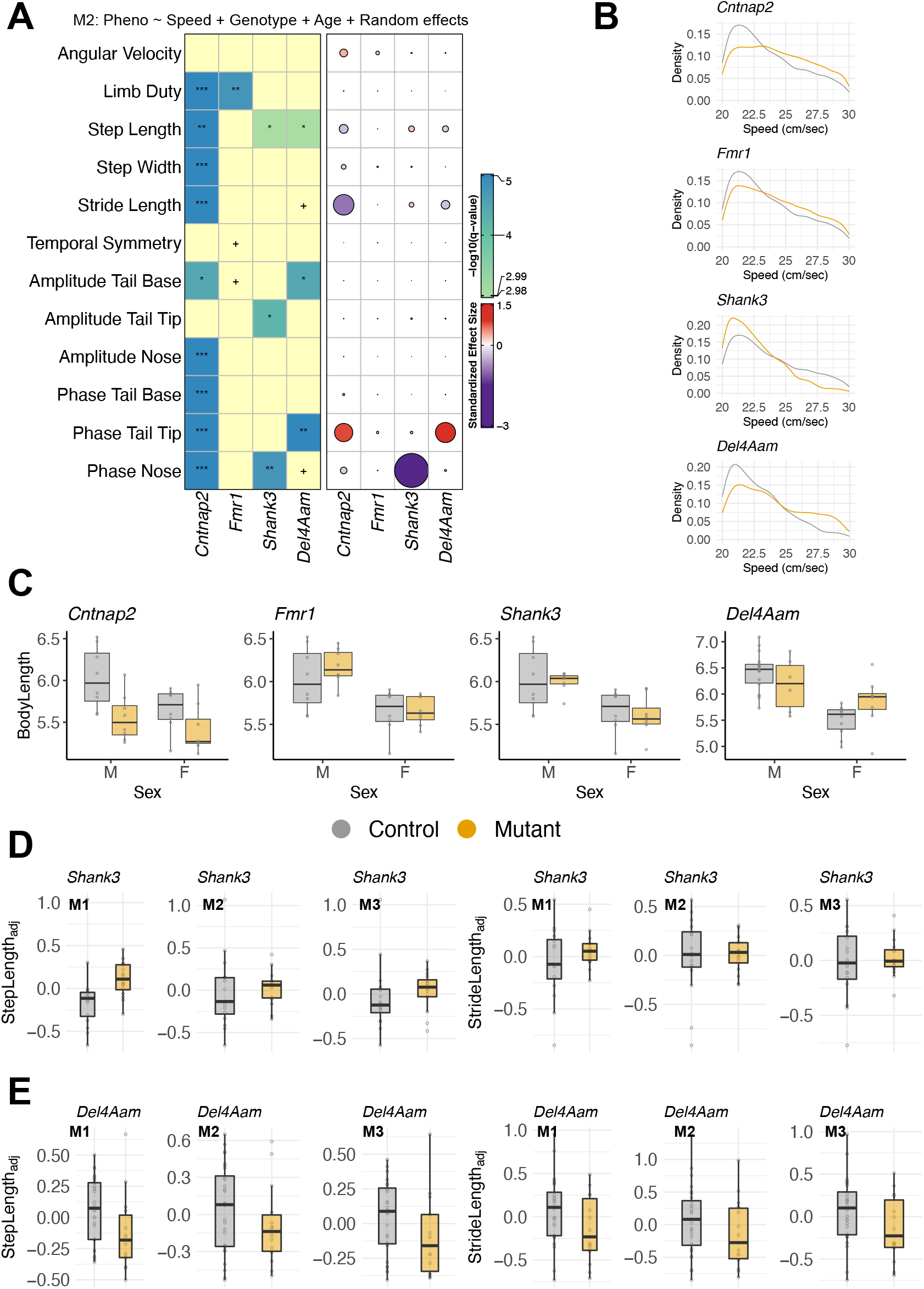
Supplemental analysis of ASD mutants described in Figure 5 (A) Heat map summarizing the effect sizes and q-values obtained from model M2: Phenotype *∼*Genotype + TestAge + Speed + (1| MouseID/TestAge). (B) Kernel density curves (estimates) of stride speed across all strains. (C) A plot showing positive association between body length and sex across different gait mutant strains. (D) Body length (M1), stride speed (M2), body length and stride speed (M3) adjusted residuals for step length and stride length for *Shank3* autism mutant. (E) Body length (M1), stride speed (M2), body length and stride speed (M3) adjusted residuals for step length and stride length for *Del4Aam* autism mutant.

**Figure S4:**
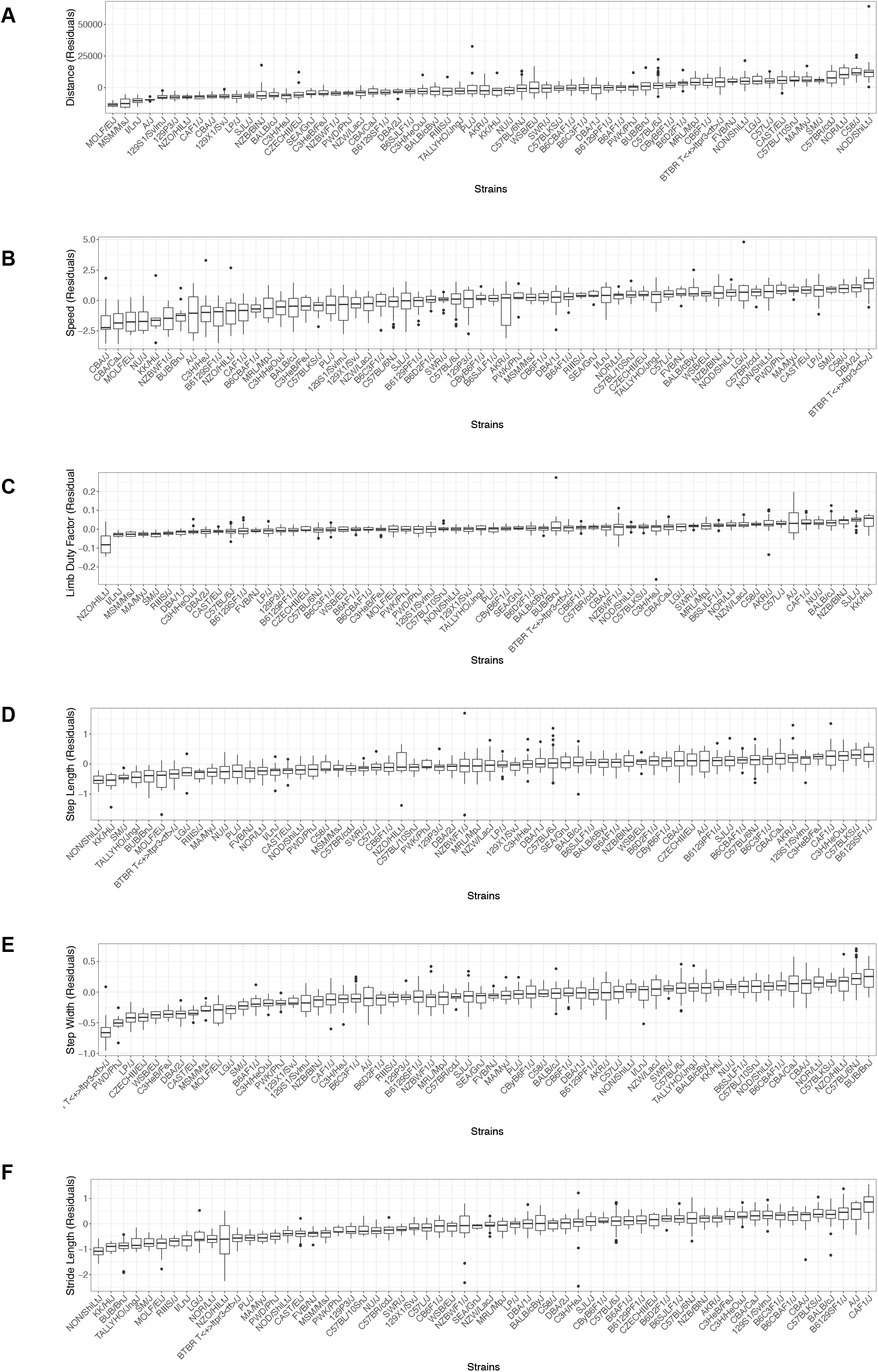
Body length adjusted phenotypes are compared across 62 strains in the strain survey. The box plots are displayed in an ascending order with respect to the median measure from left to right. Each panel (A) -(F) corresponds to a different gait phenotype.

**Figure S5:**
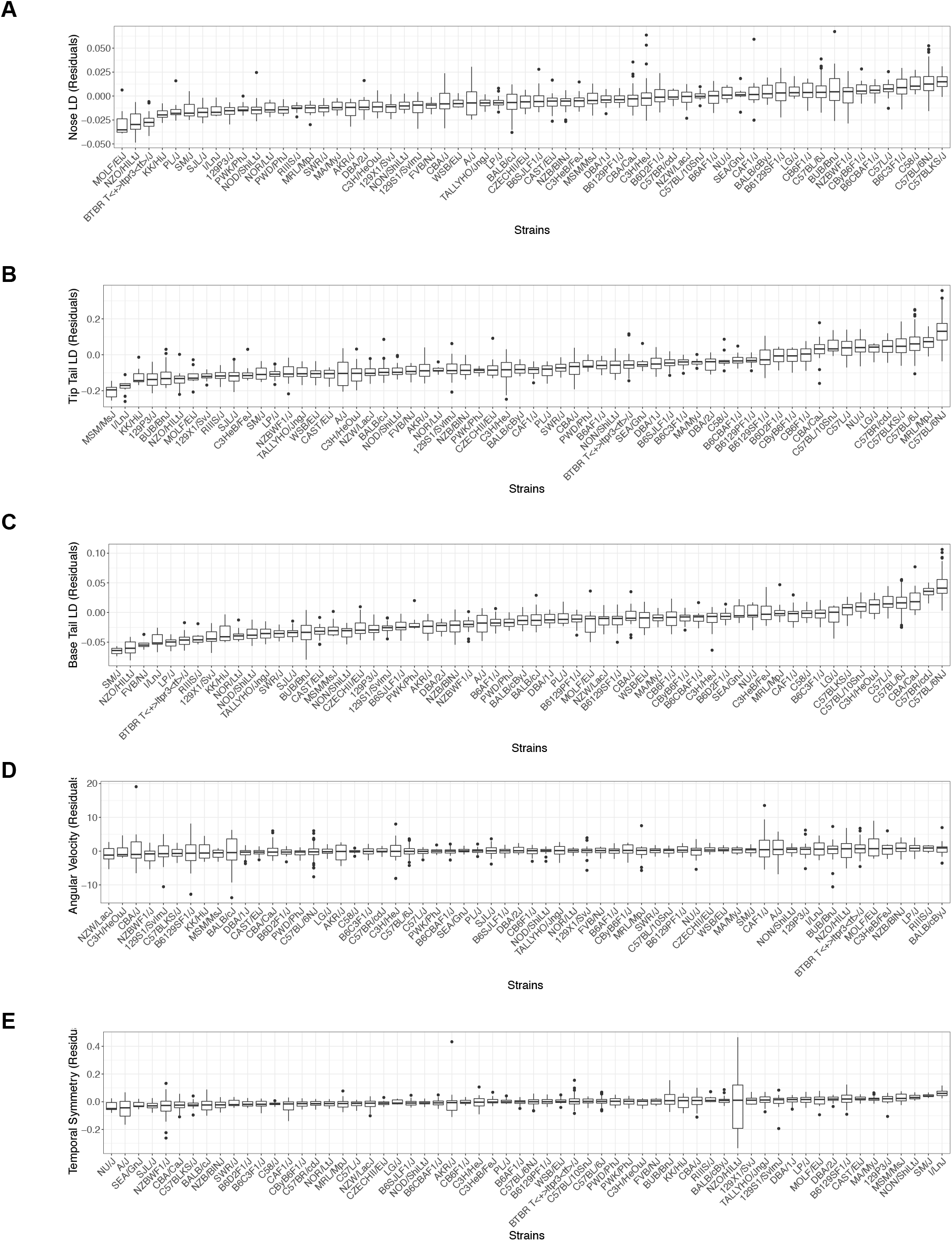
Body length adjusted phenotypes are compared across 62 strains in the strain survey. The box plots are displayed in an ascending order with respect to the median measure from left to right. Each panel (A) -(E) corresponds to a different gait phenotype.

**Figure S6:**
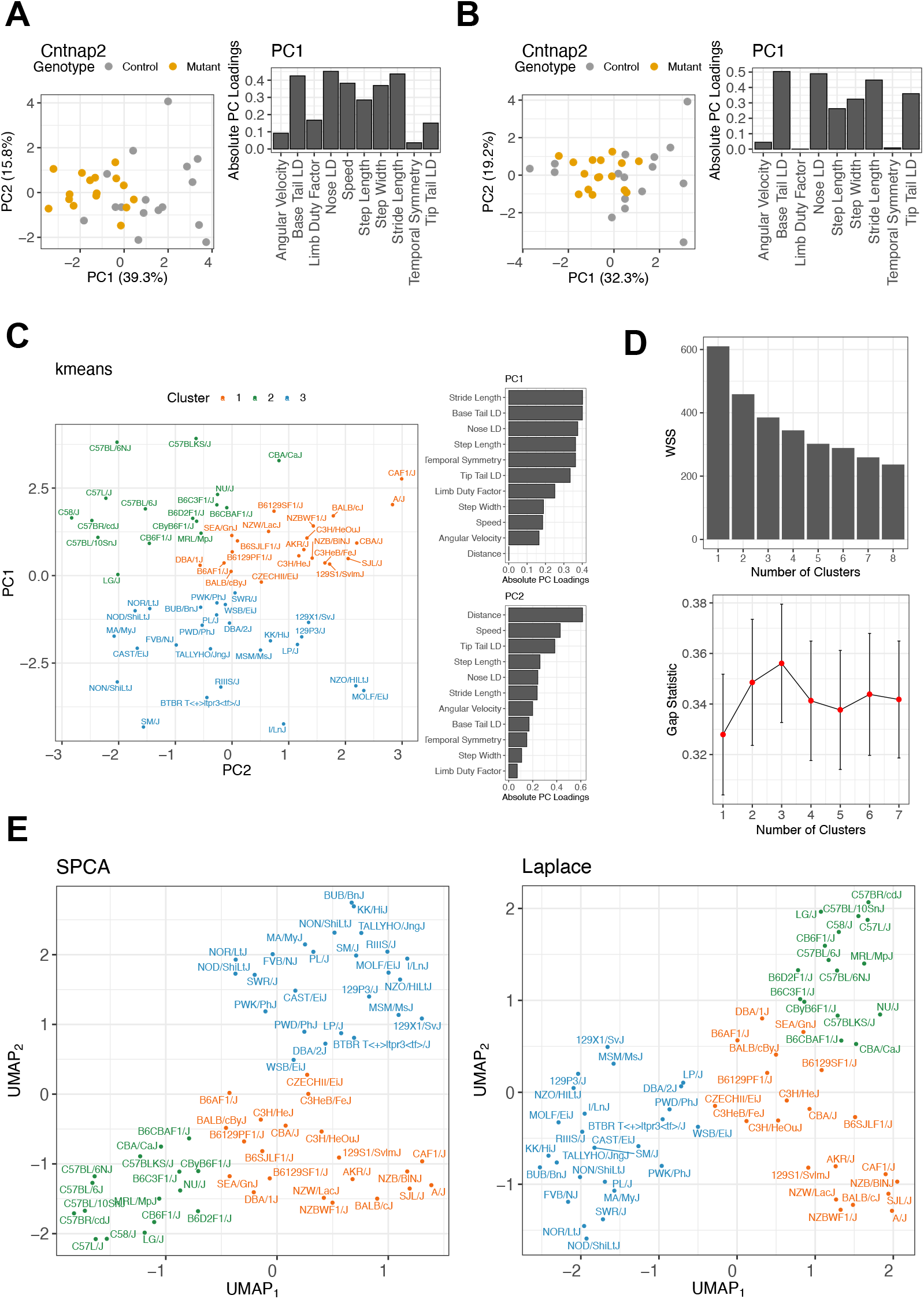
We performed PCA on z-score transformed (A) body-length adjusted gait data, (B) body-length + speed adjusted gait data. In (A), although there was some overlap between the controls and mutants along PC1, the gait phenotypes combined to separate many mutants from controls along PC1. In (B), the speed-adjusted gait phenotypes were unable to separate mutants from controls since speed was an important contributor to PC1 in (A). Removing any other important contributors to PC1 (stride length, step length, step width, nose, and base tail) by regressing them out from other phenotypes had a similar adverse effect (results not shown) on the ability to separate controls from mutants along PC1. (C) A scatterplot of the strain survey clusters with body length adjusted gait features as input is shown without the shaded areas to present a more unbiased representation of clusters in 2D PCA subspace. (D) For the k-means clusters (Figure 6C), the choice of three clusters was optimal as the gap statistic (top) [107] shows a clear peak at three clusters and the within-sum-of-squares (bottom) shows a drop at 3 clusters. (E) To see the effect of non-linear embedding, we visualized the clustering structure in a non-linear embedded space using UMAP with two different initializations (Scaled PCA, Laplace). We projected the strains to the two-dimensional UMAP space. Using the cluster memberships obtained from the k-means algorithm, we found that the UMAP dimensions preserved the separation between the three clusters discovered using the k-means.

**Figure S7:**
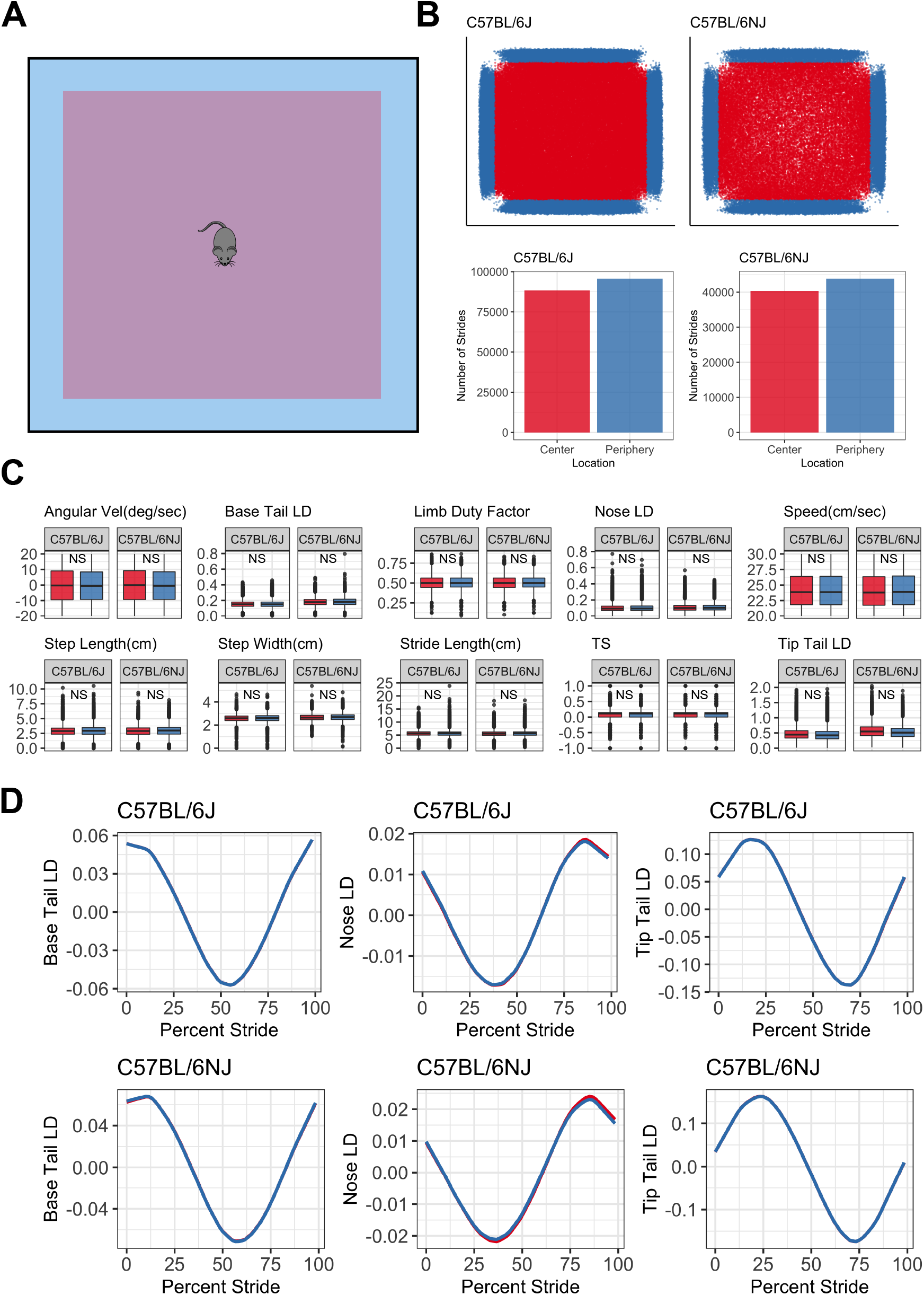
A comparative analysis of gait metrics between center and periphery of the open field arena in strains C57BL/6J and C57BL/6NJ. (A) A schematic diagram of the open field arena marks the center and periphery (outer 10%) as the central 64% area (purple) and outermost 36% area (blue). (B) Top panel shows open field with each dot representing a stride of an animal in the center (red) and periphery (blue). Bottom panel shows the number of strides in the center versus periphery for the two strains. (C) Statistical comparison of linear gait metrics between center and periphery. We fit the linear mixed model Phenotype *∼* BodyLength + TestAge + Sex + Location + (1 | MouseID) + (1| Location) where BodyLength, TestAge, Sex, Location are fixed effects and MouseID, Location are crossed random effects. To test the the null hypothesis of no effect of Location (no difference between periphery and center) on linear gait metrics, we use the *F* test with Satterthwaite’s approximation method. Our analysis revealed no significant differences across all linear gait metrics. (D) Comparison of circular gait metrics between center and periphery for the two strains showed that the phase measures are almost superimposed for base tail, nose and tip tail.

**Figure S8:**
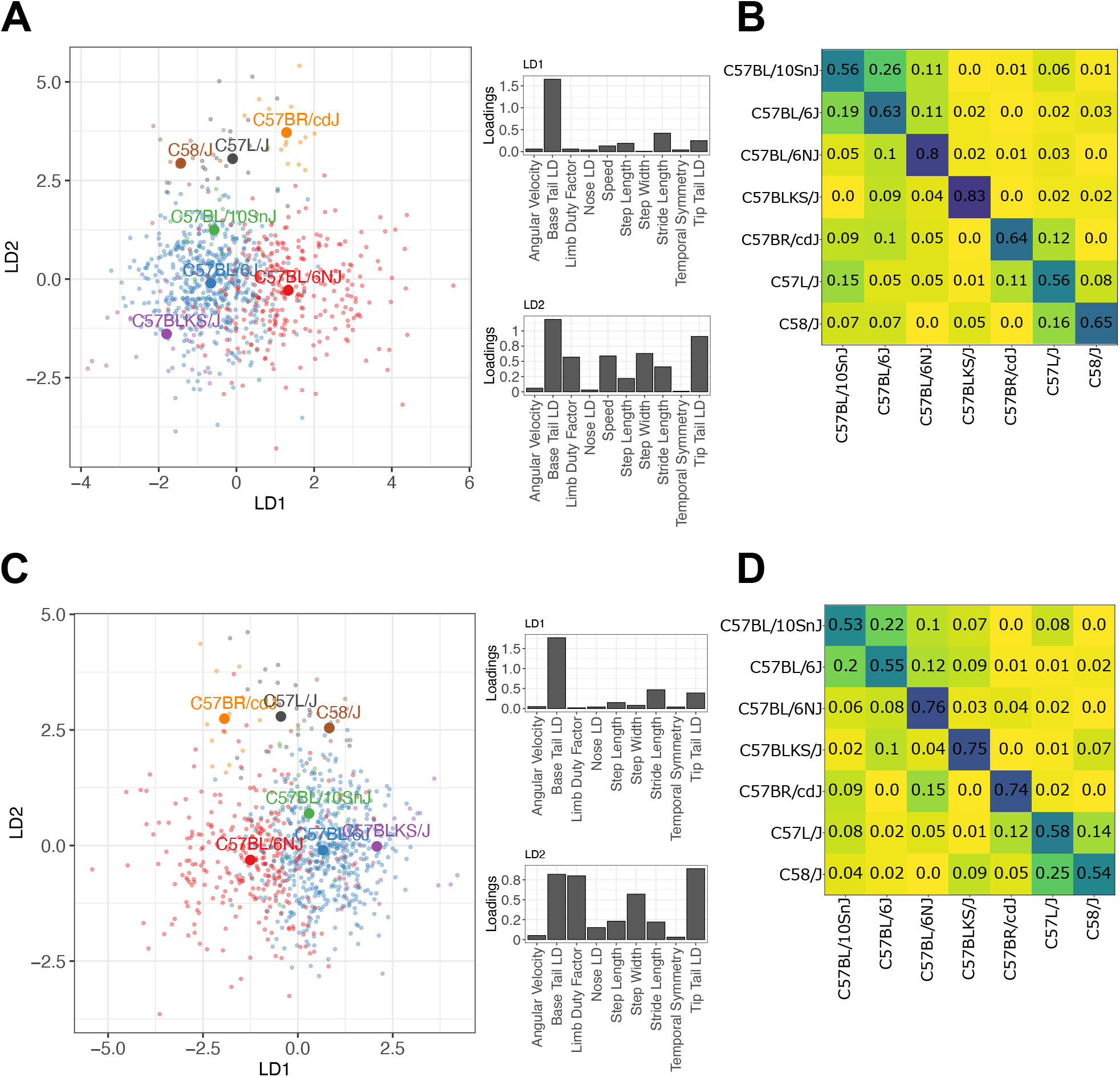
We sought to determine the gait metrics’ sensitivity to distinguish among similar strains. We took two approaches: a supervised dimension reduction approach and a multi-class classification approach. For the former, we used linear discriminant analysis [94] to quantitatively distinguish between strains C57BL/6J, C57BLKS/J, C57L/J, C57BL/6NJ, C57BR/cdJ, C57BL/10SnJ, and C58/J. First, we adjusted the gait metrics for animals’ body length by fitting a linear model with body length as a covariate. Next, we applied principal component analysis (PCA) to the residuals obtained from the linear model to address multicollinearity and prevent overfitting with LDA. We used all principal components (PCs) in the subsequent LDA algorithm to avoid the risk of throwing away critical discriminative dimensions. (A) We found that LDA separated strains when we embedded the PCs in a lower-dimensional 2D space for visualization purposes. Individual dots represent animals and dots (labeled with strain names) represent the mean/average coordinates of all animals belonging to the strain. Next, we multiplied the eigenvectors obtained from PCA with the LDA loadings matrix to identify the gait metrics that contributed to the separation between strains. For example, we found the base tail lateral displacement (Base Tail LD) to be a significant contributor to separating animals between strains C57BL/6N and C57BL/6NJ. We found similar whole-body posture differences between these two strains in our exploratory analysis earlier (see Figure 3H,I). We found the features Base Tail LD, Tip Tail LD, step width, stride speed, stride length, limb duty factor to contribute most strongly to LD2, which separated other C57 strains from C57BL/6N and C57BL/6NJ. For the second approach, we used a multi-class logistic regression (‘one versus rest’) model to predict the strain membership for each animal from its body length-adjusted gait metrics. First, we used stratified sampling to split the data into two parts: train (70%) and test (30%). Next, we used the popular resampling-based SMOTE algorithm [108] to re-balance the number of animals for each strain in the training set. We trained the classifier on the re-balanced training set and tested the performance accuracy on the test set. We performed 100 different splits on the data to allow for a proper assessment of uncertainty in our test set results. (B) We summarized our results using a normalized classification accuracy matrix that shows the proportion of correctly classified (diagonal) and misclassified (off-diagonal) animals in each strain (row). For example, for C57BL/10SnJ (first row), 56% of the test set animals were correctly classified as C57BL/10SnJ. The classifier misclassified 26% (resp. 11%) of the C57BL/10SnJ in the test set as C57BL/6J (resp. C57BL/6NJ). (C,D) We performed similar analyses as in (A,B) except the gait metrics were not adjusted for both body length and stride speed of the animals.

**Figure S9:**
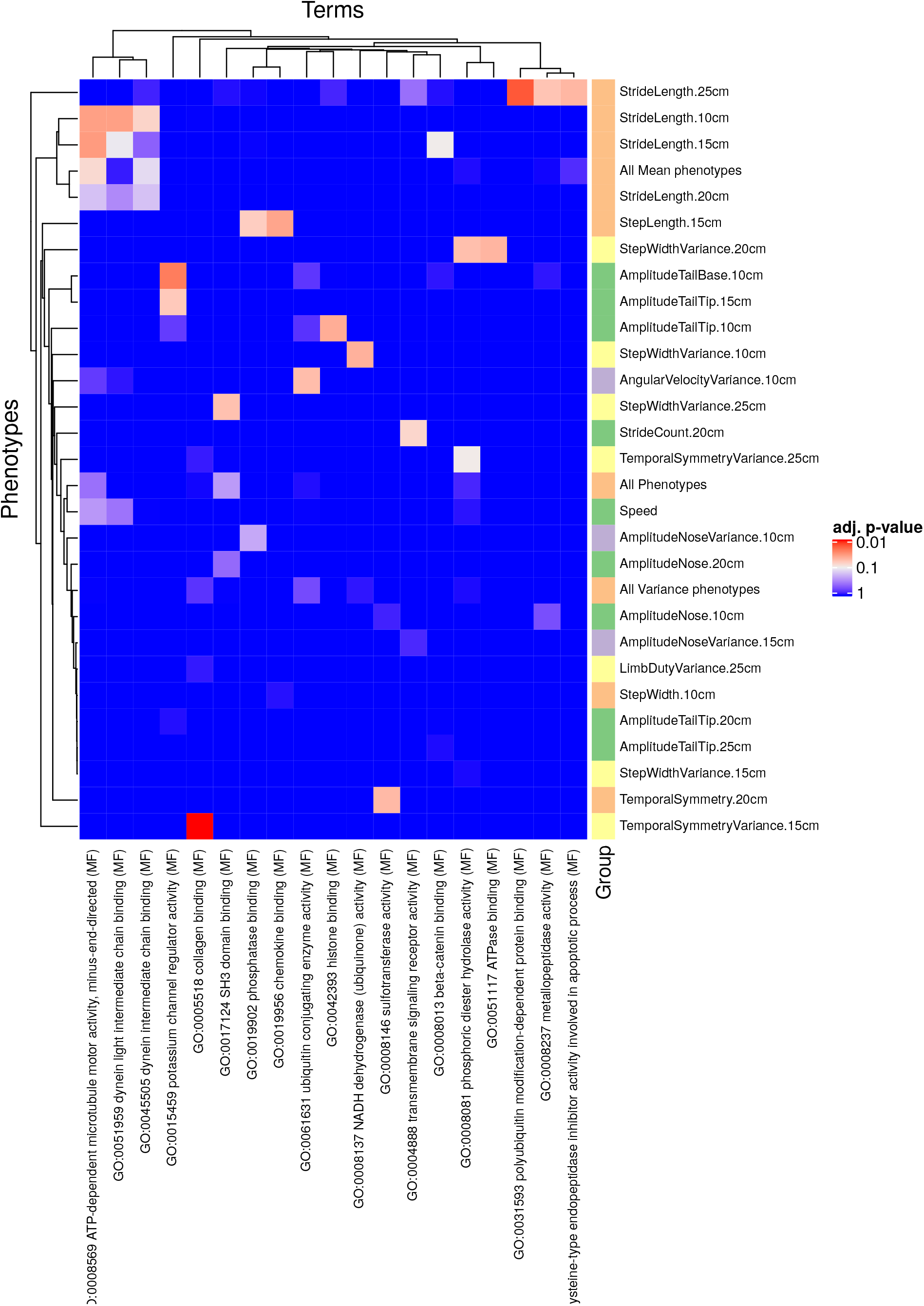
INRICH results for GO terms, KEGG pathways and mouse phenotypes. The heatmaps contain terms and phenotypes that passed the 0.1 corrected p-value threshold.

**Table S3:** A table summarizing effect sizes and q-values (FDR-adjusted p-values, padj) obtained from models M1,M2,M3 for all phenotypes and gait strains.

| M1 : Phenotype ~ Genotype + TestAge + BodyLength + (1 \| MouseID/TestAge) |  |  |  |  |  |  |  |  |
| --- | --- | --- | --- | --- | --- | --- | --- | --- |
| Phenotype | Sod1 |  | Ts65Dn |  | Mecp2 (Het) |  | Mecp2 (Hemi) |  |
|  | Effect size | P<sub>adj</sub> | Effect size | P<sub>adj</sub> | Effect size | P<sub>adj</sub> | Effect size | P<sub>adj</sub> |
| Angular Velocity | 0.02 | 0.69 | 0.04 | 0.69 | 0.02 | 0.69 | -0.25 | 0.37 |
| Speed | -0.21 | 0.00 | 0.15 | 0.04 | -0.38 | 0.00 | -0.53 | 0.00 |
| Limb Duty | 0.23 | 0.00 | -0.10 | 0.21 | 0.03 | 0.74 | 0.27 | 0.18 |
| Step Length | -0.12 | 0.34 | -0.24 | 0.02 | -0.10 | 0.37 | -0.38 | 0.28 |
| Step Width | -0.09 | 0.56 | 0.59 | 0.00 | 0.16 | 0.48 | -0.79 | 0.02 |
| Stride Length | -0.30 | 0.01 | -0.35 | 0.01 | -0.33 | 0.06 | -0.60 | 0.01 |
| Temporal Symmetry | -0.09 | 0.24 | 0.12 | 0.15 | -0.00 | 0.96 | 0.09 | 0.64 |
| Amplitude Tail Base | 0.23 | 0.11 | -0.46 | 0.03 | 0.34 | 0.11 | 0.40 | 0.09 |
| Amplitude Tail Tip | 0.16 | 0.12 | -0.37 | 0.00 | 0.50 | 0.01 | 0.99 | 0.00 |
| Amplitude Nose | 0.02 | 0.73 | -0.16 | 0.16 | 0.09 | 0.53 | 0.09 | 0.53 |
| Phase Tail Base | 0.02 | 0.11 | -0.05 | 0.00 | 0.00 | 0.39 | 0.04 | 0.01 |
| Phase Tail Tip | 0.09 | 0.16 | -1.37 | 0.00 | -0.17 | 0.02 | 0.50 | 0.00 |
| Phase Nose | -0.06 | 0.01 | -0.64 | 0.00 | -0.05 | 0.06 | -0.18 | 0.00 |
| M2 : Phenotype ~ Genotype + TestAge + Speed + (1 \| MouseID/TestAge) |  |  |  |  |  |  |  |  |
| Angular Velocity | 0.02 | 0.68 | 0.03 | 0.68 | 0.02 | 0.68 | -0.14 | 0.28 |
| Limb Duty | 0.14 | 0.12 | -0.03 | 0.71 | -0.10 | 0.36 | -0.23 | 0.02 |
| Step Length | -0.08 | 0.66 | -0.23 | 0.15 | 0.03 | 0.83 | -0.77 | 0.00 |
| Step Width | -0.12 | 0.59 | 0.59 | 0.00 | 0.14 | 0.59 | -1.12 | 0.00 |
| Stride Length | -0.24 | 0.16 | -0.39 | 0.02 | -0.11 | 0.59 | -1.09 | 0.00 |
| Temporal Symmetry | -0.04 | 0.65 | 0.08 | 0.45 | 0.04 | 0.65 | 0.34 | 0.00 |
| Amplitude Tail Base | 0.21 | 0.15 | -0.45 | 0.03 | 0.28 | 0.15 | 0.30 | 0.15 |
| Amplitude Tail Tip | 0.13 | 0.17 | -0.37 | 0.00 | 0.46 | 0.01 | 0.90 | 0.00 |
| Amplitude Nose | -0.00 | 0.95 | -0.14 | 0.21 | 0.05 | 0.87 | 0.04 | 0.87 |
| Phase Tail Base | 0.01 | 0.22 | -0.05 | 0.00 | 0.01 | 0.34 | 0.00 | 0.08 |
| Phase Tail Tip | 0.00 | 0.29 | -4.55* | 0.01 | -10.50* | 0.06 | 0.28 | 0.00 |
| Phase Nose | -0.08 | 0.05 | -8.14* | 0.00 | -0.09 | 0.13 | -4.12* | 0.00 |
| M3 : Phenotype ~ Genotype + TestAge + Speed + BodyLength + (1 \| MouseID/TestAge) |  |  |  |  |  |  |  |  |
| Angular Velocity | 0.02 | 0.67 | 0.03 | 0.67 | 0.02 | 0.67 | -0.24 | 0.42 |
| Limb Duty | 0.14 | 0.13 | -0.03 | 0.79 | -0.12 | 0.26 | 0.04 | 0.79 |
| Step Length | -0.07 | 0.65 | -0.27 | 0.01 | -0.01 | 0.92 | -0.28 | 0.55 |
| Step Width | -0.11 | 0.49 | 0.60 | 0.00 | 0.12 | 0.49 | -0.81 | 0.02 |
| Stride Length | -0.22 | 0.07 | -0.40 | 0.00 | -0.18 | 0.32 | -0.41 | 0.07 |
| Temporal Symmetry | -0.05 | 0.41 | 0.09 | 0.31 | 0.06 | 0.41 | 0.18 | 0.31 |
| Amplitude Tail Base | 0.21 | 0.19 | -0.45 | 0.03 | 0.28 | 0.19 | -0.35 | 0.23 |
| Amplitude Tail Tip | 0.13 | 0.17 | -0.37 | 0.00 | 0.46 | 0.01 | 0.90 | 0.00 |
| Amplitude Nose | -0.00 | 0.95 | -0.14 | 0.21 | 0.05 | 0.87 | 0.04 | 0.87 |
| Phase Tail Base | 0.01 | 0.34 | -0.05 | 0.00 | 0.01 | 0.34 | 0.00 | 0.34 |
| Phase Tail Tip | 0.00 | 0.03 | -2.96* | 0.00 | -0.71 | 0.22 | 0.21 | 0.01 |
| Phase Nose | -0.08 | 0.05 | -5.60* | 0.00 | -275.28* | 0.22 | -8.24* | 0.00 |

**Table S4:** A table summarizing effect sizes and q-values (FDR-adjusted p-values, padj) obtained from models M1,M2,M3 for all phenotypes and autism strains.

| M1 : Phenotype ~ Genotype + BodyLength + Sex + (1 \| MouseID/TestAge) |  |  |  |  |  |  |  |  |
| --- | --- | --- | --- | --- | --- | --- | --- | --- |
| Phenotype | *Cntnap2* |  | *Fmr1* |  | *Shank3* |  | *Del4Aam* |  |
|  | Effect size | P<sub>adj</sub> | Effect size | P<sub>adj</sub> | Effect size | P<sub>adj</sub> | Effect size | P<sub>adj</sub> |
| Angular Velocity | 0.51 | 0.79 | 0.13 | 0.83 | 0.13 | 0.83 | -0.17 | 0.83 |
| Speed | 1.07 | 0.00 | 0.64 | 0.00 | -0.90 | 0.00 | 0.77 | 0.02 |
| Limb Duty | 0.02 | 0.00 | 0.01 | 0.10 | 0.00 | 0.84 | -0.01 | 0.41 |
| Step Length | -0.19 | 0.13 | 0.03 | 0.77 | 0.23 | 0.08 | -0.18 | 0.08 |
| Step Width | -0.17 | 0.01 | -0.08 | 0.15 | -0.04 | 0.50 | -0.01 | 0.90 |
| Stride Length | -0.59 | 0.00 | 0.04 | 0.68 | 0.13 | 0.25 | -0.23 | 0.08 |
| Temporal Symmetry | 0.00 | 0.66 | -0.01 | 0.21 | -0.01 | 0.21 | -0.00 | 0.87 |
| Amplitude Tail Base | -0.01 | 0.01 | -0.01 | 0.08 | -0.00 | 0.45 | -0.02 | 0.01 |
| Amplitude Tail Tip | 0.02 | 0.29 | -0.03 | 0.27 | -0.06 | 0.02 | -0.04 | 0.14 |
| Amplitude Nose | -0.02 | 0.00 | -0.01 | 0.07 | -0.00 | 0.49 | -0.01 | 0.07 |
| Phase Tail Base | 0.07 | 0.00 | 0.02 | 0.19 | 0.00 | 0.49 | -0.02 | 0.24 |
| Phase Tail Tip | 0.71 | 0.00 | -0.03 | 0.39 | -0.12 | 0.15 | 0.52 | 0.00 |
| Phase Nose | -0.09 | 0.00 | -0.01 | 0.28 | -0.12 | 0.00 | -0.11 | 0.04 |
| M2 : Phenotype ~ Genotype + Speed + (1 \| MouseID/TestAge) |  |  |  |  |  |  |  |  |
| Angular Velocity | 0.35 | 0.96 | 0.16 | 0.96 | 0.05 | 0.96 | -0.04 | 0.96 |
| Limb Duty | 0.03 | 0.00 | 0.02 | 0.01 | -0.01 | 0.12 | 0.00 | 0.85 |
| Step Length | -0.40 | 0.00 | 0.01 | 0.91 | 0.26 | 0.05 | -0.27 | 0.05 |
| Step Width | -0.21 | 0.00 | -0.06 | 0.36 | -0.05 | 0.36 | -0.01 | 0.87 |
| Stride Length | -0.91 | 0.00 | -0.01 | 0.94 | 0.21 | 0.11 | -0.38 | 0.08 |
| Temporal Symmetry | 0.00 | 0.66 | -0.02 | 0.09 | -0.01 | 0.66 | -0.01 | 0.66 |
| Amplitude Tail Base | -0.01 | 0.01 | -0.01 | 0.10 | -0.00 | 0.36 | -0.02 | 0.01 |
| Amplitude Tail Tip | 0.02 | 0.24 | -0.03 | 0.24 | -0.07 | 0.01 | -0.04 | 0.13 |
| Amplitude Nose | -0.02 | 0.00 | -0.00 | 0.14 | -0.00 | 0.23 | -0.01 | 0.14 |
| Phase Tail Base | 0.09 | 0.00 | 0.02 | 0.19 | -0.00 | 0.48 | -0.02 | 0.26 |
| Phase Tail Tip | 0.81 | 0.00 | -0.10 | 0.14 | -0.11 | 0.14 | 0.89 | 0.00 |
| Phase Nose | -0.30 | 0.00 | -0.03 | 0.16 | -0.21 | 0.01 | -0.09 | 0.07 |
| M3 : Phenotype ~ Genotype + Speed + BodyLength + (1 \| MouseID/TestAge) |  |  |  |  |  |  |  |  |
| Angular Velocity | 0.54 | 0.68 | 0.11 | 0.98 | 0.05 | 0.98 | 0.02 | 0.98 |
| Limb Duty | 0.03 | 0.00 | 0.02 | 0.01 | -0.01 | 0.16 | 0.00 | 0.78 |
| Step Length | -0.26 | 0.04 | -0.01 | 0.91 | 0.28 | 0.03 | -0.24 | 0.03 |
| Step Width | -0.17 | 0.01 | -0.08 | 0.17 | -0.04 | 0.39 | 0.00 | 0.98 |
| Stride Length | -0.73 | 0.00 | -0.05 | 0.66 | 0.24 | 0.03 | -0.32 | 0.02 |
| Temporal Symmetry | -0.00 | 0.59 | -0.02 | 0.11 | -0.01 | 0.59 | -0.01 | 0.59 |
| Amplitude Tail Base | -0.02 | 0.00 | -0.01 | 0.11 | -0.01 | 0.34 | -0.02 | 0.00 |
| Amplitude Tail Tip | 0.02 | 0.24 | -0.03 | 0.24 | -0.07 | 0.01 | -0.04 | 0.13 |
| Amplitude Nose | -0.02 | 0.00 | -0.00 | 0.14 | -0.00 | 0.23 | -0.01 | 0.14 |
| Phase Tail Base | 0.09 | 0.00 | 0.02 | 0.19 | 0.00 | 0.47 | -0.02 | 0.26 |
| Phase Tail Tip | 0.31 | 0.00 | -0.29 | 0.01 | 0.07 | 0.35 | 0.39 | 0.00 |
| Phase Nose | -0.38 | 0.00 | -0.03 | 0.16 | -225.04 | 0.02 | -0.13 | 0.08 |

